# Bioinformatic Analysis Linking Genomic Defects to Chemosensitivity and Mechanism of Action

**DOI:** 10.1101/2020.11.20.391029

**Authors:** David G. Covell

## Abstract

A joint analysis of NCI60 small molecule screening data, their genetically defective genes and mechanisms of action (MOA) of FDA approved cancer drugs screened in the NCI60 is proposed for identifying links between chemosensitivity, genomic defects and MOA. Self-organizing-maps (SOMs) are used to organize the chemosensitivity data. Student’s t-tests are used to identify SOM clusters with chemosensitivity for tumor cells harboring genetically defective genes. Fisher’s exact tests are used to reveal instances where defective gene to chemosensitivity associations have enriched MOAs. The results of this analysis find a relatively small set of defective genes, inclusive of ABL1, AXL, BRAF, CDC25A, CDKN2A, IGF1R, KRAS, MECOM, MMP1, MYC, NOTCH1, NRAS, PIK3CG, PTK2, RPTOR, SPTBN1, STAT2, TNKS and ZHX2, as possible candidates for roles in chemosensitivity for compound MOAs that target primarily, but not exclusively, kinases, nucleic acid synthesis, protein synthesis, apoptosis and tubulin. This analysis may contribute towards the goals of cancer drug discovery, development decision making, and explanation of mechanisms.

## INTRODUCTION

The emergence of extensive human tumor cell line compound screening data, coupled with advances in cancer genomic technologies, has generated comprehensive and complex databases [1]. Strategies for analyzing this data may provide important links between genetic changes that contribute to the hallmarks of cancer biology [2] and the discovery of leads in the pursuit of small-molecule cancer therapy [3]. The present report examines links between genetically defective genes in the NCI60 tumor cells, chemosensitivity, as measured by growth inhibition (<u>GI50</u>_NCI60;_ adopting the convention of an <u>under bar</u> to describe the vector of <u>GI50</u>_NCI60_ (N=59) measurements for each screened compound) and preferences for mechanisms of action (MOA) of identified linkages. An elegant study by Ikediobi et al. [4] addressed this goal by examining relationships between mutations in 24 cancer genes in the NCI60 tumor cells and the <u>GI50</u>_NCI60_ activity of ~8k screened compounds. Their finding of a strong association between the BRAF mutation (V600E) and the <u>GI50</u>_NCI60_ activity of phenothiazines supports important links between defective genes, chemosensitivity and MOAs. The current analysis extends this work, with significant differences;

- <u>GI50</u>_NCI60_ results for ~53k screened compounds were analyzed (DTP database).
- A larger set of gene mutations (N=365) for the NCI60 tumor cell lines was analyzed (CBioPortal database).
- A novel analysis of <u>GI50</u>_NCI60_, based on Self-Organizing-Maps (SOMs), emphasizing FDA approved compounds with assigned MOAs in the NCI60 screened compounds, was used to derive links between tumor cell chemosensitivity, genetically defective genes and MOAs.

## METHODS/DATA

A flow chart describing the data to be analyzed and the methods of analysis appears in **Figure 1**. Each of the five boxes in this figure (labelled A through E) will be discussed. **Box A** describes the NCI60 Data and clustering procedures. Historically the NCI60 screen was designed to identify relationships between chemotypes and cellular responses [5]. Many statistical tools are now available for clustering <u>GI50</u>_NCI60_ data [6]. Relying on our prior analyses [7], the results presented here use self-organizing-maps (SOMs) to cluster <u>GI50</u>_NCI60_ data. The currently available DTP data consists of ~53k screened compounds, which for this analysis was reduced to 46,798 data records when filtered for a coefficient of variation above 0.1. Parameters from prior SOM analyses were selected for clustering (hexagonal nearest neighbors, EP kernel[8]). SOM dimensions, based on a heuristic using the ratio of the first two principal components of <u>GI50</u>_NCI60_ vectors, yielded map dimensions with 44 rows and 28 columns (referred to hereafter as SOM_DTP_). Each of these 1232 SOM nodes yields a vector representing the ‘average’ <u>GI50</u>_NCI60_ for all compounds clustered within each SOM node (referred to hereafter as a node’s <u>GI50</u>_codebook_). Reference to SOM_DTP_ nodes represents a cluster of similar <u>GI50</u>_NCI60_ values. The appearance of a screened compound in a SOM_DTP_ node will be referred to as a its SOM ‘projection’. Prior analyses find <u>GI50</u>_codebook_ patterns to be associated with a compound’s MOA (e.g. alkylating agents, tubulin targeting agents, DNA/RNA damaging agents, agents affecting mitochondrial function [7]). Analysis of SOM-derived <u>GI50</u>_codebook_ patterns has also been proposed for use in the development of clinical strategies based on differentially expressed molecular targets within classes of tumors [9, 10]. For example, the recent identification of unique <u>GI50</u>_codebook_ patterns within the NCI60 renal subpanel has provided the basis for further testing of the natural product-derived family of englerins [11].

**Figure 1:**
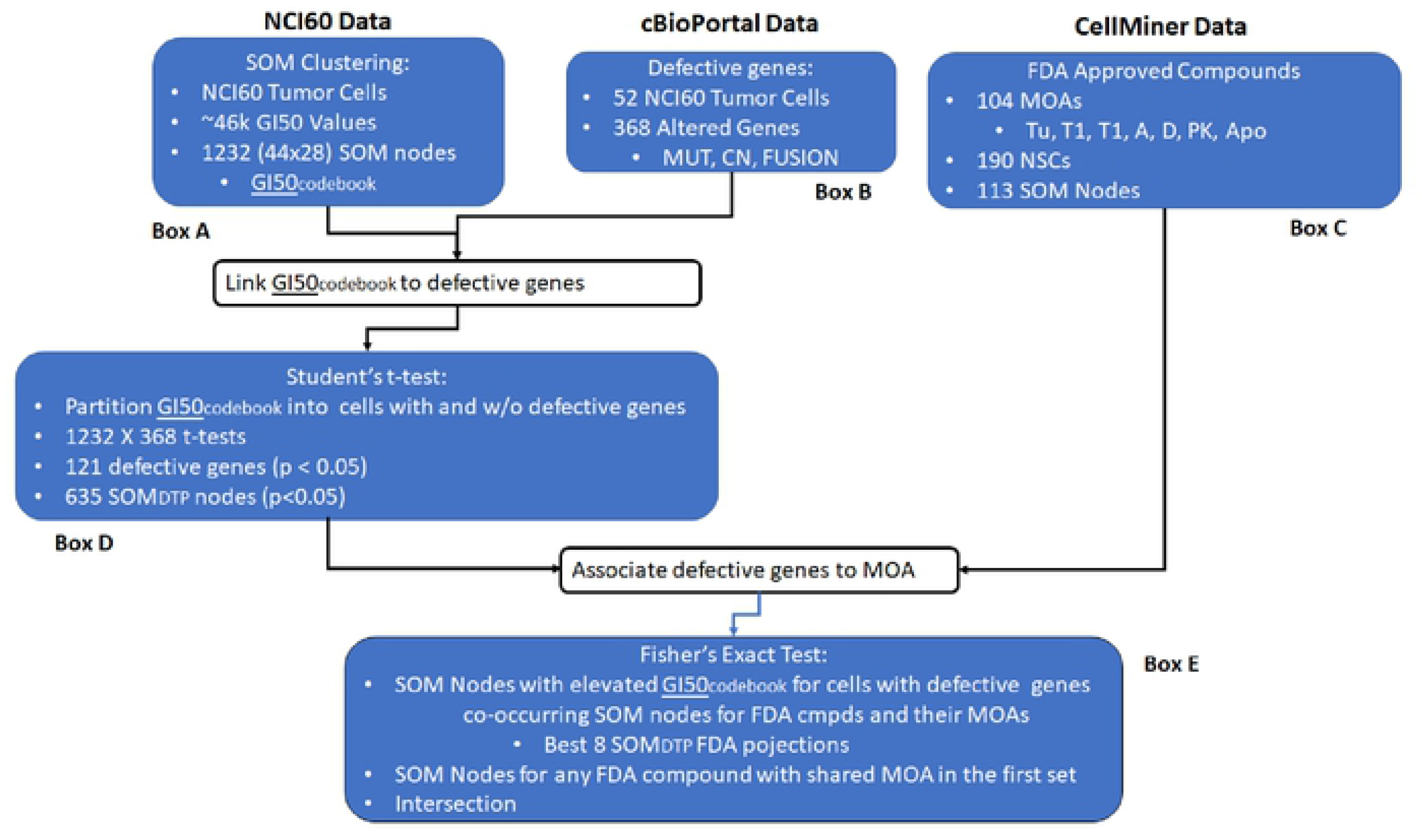
Flow chartfor the proposed analysis. See text for descriptions of **Boxes A-E**.

**Box B** describes the defective gene data within the NCI60. The cBioPortal database https://www.cbioportal.org/ [12, 13] lists a total of 368 genes in the NCI60 that have defective genes; as either a mutation (MUT), copy number alteration (CNA) or fusion/splice (FUSION). The analysis reported here groups all genomic changes, so that any genes designated as genetically defective will be due to any or all variations. Defective genes occur within each NCI60 tumor cell individually or as pairs, doublets, triplets, etc. **Appendix Figure 1** displays a histogram for the frequency of defective genes within the NCI60. The highest frequency exists for tumor cells having a single defective gene. This frequency decreases progressively down to less than one percent for tumor cells sharing 10 defective genes. The cumulative frequency of tumor cells sharing any defective gene is 0.97, an indication that the probability of tumor cells sharing any defective gene is approaches one. **Appendix Figure 2** displays the histogram of defective genes shared between tumor cells. These results find that shared defective genes, comprised of doublets and triplets of defective genes are more common (higher frequency) compared to the appearance of only a single defective gene (consistent with Ikediobi et al.[4]). **Appendix Table I** lists the singlets, doublets and triplets of defective genes observed in the NCI60. **Appendix Table II** summarizes their counts. Inspection finds CDC25A, TP53, CDKN2A, CDKN2B, MYC, BRAF, EP300, KRAS, NOTCH1 and PTK2 as the top ten most frequently occurring defective genes. To summarize, defective genes appearing as doublets or triplets finds these top ten defective genes to appear in combination with themselves and other genes. Collectively these results indicate that shared defective genes are common within the NCI60.

**Box C** provides information about MOAs for FDA approved compounds. CellMiner [14] reports 270 FDA compounds with unique NSC and Name assignments that have been screened in the NCI60 (ca. 2019). One-hundred and ninety FDA screened compounds appear in the 46,798 <u>GI50</u>_NCI50_ responses; filtered for a coefficient of variation above 0.1. One-hundred and four MOA assignments exist for this set of compounds. These assignments consist of a primary MOA designation followed by secondary MOAs. For example the assignment BCR-ABL|YK,FYN,LYN indicates BCR-ABL at the primary MOA, with YK (tyrosine kinase), FYN and LYN as secondary MOA assignments. The complete set of MOAs for FDA screened compounds appears as **Appendix Table III**. Seven primary MOA classes spanning this data function to target tubulin:**Tu**, topoisomerase 2:**T2**, topoisomerase 1:**T1**, alkylation:**A** (A2: Alkylating_at_N-2_position_of_guanine, A6: Alkylating_at_O-6_of_guanine, A7: Alkylating_at_N-7_position_of_guanine, AlkAg: Alkylating agent and anti-metabolites:AM), DNA:**D** (Db:DNA_binder, DDI/R, DNA_damage_repair/inducer, Df: antifols, Ds: DNA_synthesis_inhibitor), kinases:**PK** and apoptosis:**Apo**. MOA:**PK** consists of over 100 kinase targets. FDA compounds screened in the NCI60, and their assigned CellMiner MOA, are projected to 113 of the 1232 SOM_DTP_ nodes. The intersection of these nodes with SOM_DTP_ nodes having relatively greater chemosensitivity for tumor cells with and without defective genes will be reported. A previous study of FDA approved compounds screened in the NCI60, using correlative <u>GI50</u>_NCI60_ measures, has been reported by Holbeck et al. Their emphasis only on drugs that target signal transduction represents an overlapping subset of the data presented here.

**Box D** describes the steps for linking chemosensitivity, defined by <u>GI50</u>_codebook_, to defective genes. The process begins by partitioning each <u>GI50</u>_codebook_ into subsets comprising tumor cells with (e.g. <u>GI50</u>_defective_) and without (<u>GI50</u>_wt_) a defective gene and applying a Student’s t-test to identify cases of relatively higher chemosensitivity for tumor cells with defective genes. SOM_DTP_ nodes with Student’s p-values below 0.05 were further assessed for statistical significance by resampling. Each node’s <u>GI50</u>_codebook_ was randomly shuffled and the Student’s t-test performed, while maintaining the tumor cell’s wt and defective gene status. One-thousand trials were conducted for each <u>GI50</u>_codebook_ and a p-value was estimated by counting the number of times the shuffled p-value was less than the initial, unshuffled, p-value. Dividing this value by 1000 yields an estimate for the probability of the observed p-value occurring by chance. SOM nodes with measured p-values less than 0.05 and below their estimated chance occurrence were accepted for further analysis. Sixty-five percent (65%, n=635) of the 1232 SOM nodes pass this criterion and account for 121 defective genes.

**Box E** describes the process of associating MOAs and defective genes using a Fisher’s exact test. This test is useful for categorical data that result from classifying objects in two different ways; it is used to examine the significance of the association (contingency) between these two kinds of classification. Each FDA approved agent and its assigned MOA that has been screened in the NCI60 is associated with the SOM node that best matches the compound’s <u>GI50</u>_NCI60_ to a SOM_DTP_ node’s <u>GI50</u>_codebook_. (cf. **RESULTS Figure 2**, blue hexagons). This association can be extended to identify the 2^nd^, 3^rd^, 4^th^, etc. best matching SOM projections, thereby establishing the SOM region (comprised of multiple nodes) representing the cluster location of an FDA approved compound and its MOA. Typically, the best matching SOM_DTP_ node is all that may be needed [7], however extending the SOM_DTP_ region beyond only the best node establishes greater confidence in a SOM_DTP_ region for projecting an FDA compound. Fisher’s exact statistics will use the best eight SOM_DTP_ projections for FDA approved compounds. Fisher’s exact contingency tables are based on two data stratifications. First, for each defective gene, SOM_DTP_ nodes for FDA approved compounds are collected from only nodes having significantly higher chemosensitivity for <u>GI50</u>_defective_ compared to <u>GI50</u>_wt_. The second stratification selects SOM_DTP_ nodes for all FDA compounds that share the MOA of any compound in the first classification, ignoring whether the SOM_DTP_ node has relatively greater chemosensitivty for tumor cells with the defective gene. The contingency table, for input to the Fisher’s exact test, consists of the SOM_DTP_ nodes for MOA’s of FDA compounds that also share relatively greater chemosensitivity for tumor cells with defective genes, SOM_DTP_ nodes for any FDA compound that shares any MOA in the first set and their intersection. The Fisher’s exact score is a statistical measure for the random likelihood of the intersection of nodes from each classification. Results are reported using the log(pvalue) for each Fisher’s exact test.

**Figure 2:**
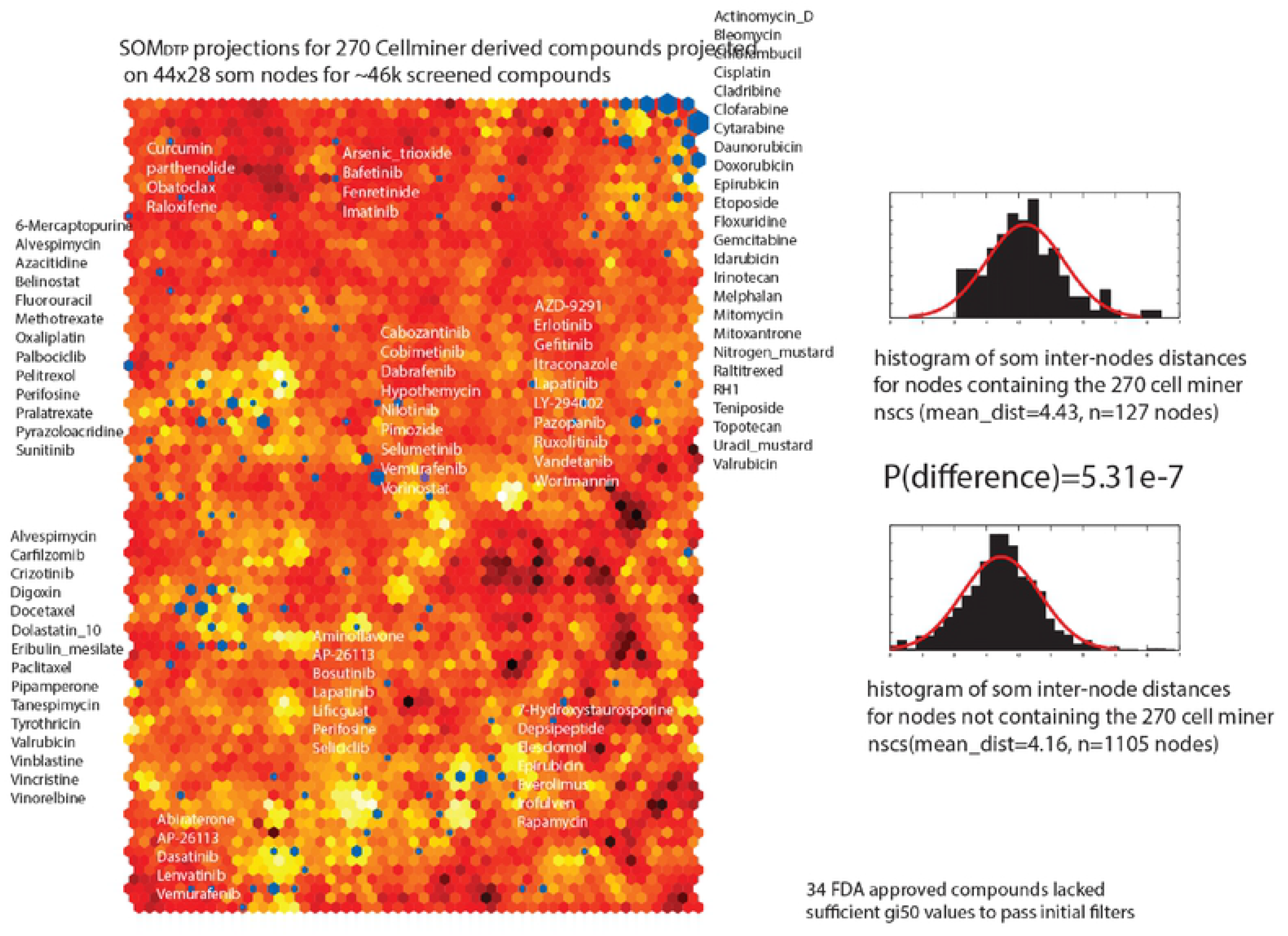
SOM_DTP_ is colored according similarity of <u>GI50</u>_codebooks_, where the most similar node neighbors are displayed in deep red and the most disi-similar node neighbors appear in bright yellow. FDA approved compounds are projected onto SOM_DTP_ as blue hexagons, where hexagons are sized according to the number of FDA agents appearing in any node. FDA compound names are displayed adjacent to their SOM_DTP_ projections. FDA approved projections to SOM_DTP_ nodes are listed in Appendix Table III.

## RESULTS

The results from **Box A** in the flow chart reveal important features about the SOM analysis of <u>GI50</u>_NCI60_ and the cluster locations for FDA approved compounds. SOM analysis maps the <u>GI50</u>_NCI60_ data into a 2-dimensional image of 1232 nodes (44X28), referred to above as SOM_DTP_. Each SOM_DTP_ node consists of <u>GI50</u>_NCI60_ values that cluster together; represented by an average vector (SOM_codebook_, as discussed above). SOMs are further organized by placing similar SOM_codebooks_ as nearest neighbors; a distinguishing feature of SOMs which, in this case, assigns a node’s position according to similarity with its six (i.e. hexagonal) neighbors. This feature is distinctive from typical hierarchical clustering which define clade neighbors according to pair-wise comparisons.

**Figure 2** displays, in the left panel, SOM_DTP_, colored according to similarity of neighboring <u>GI50</u>_codebooks_; where the most similar <u>GI50</u>_codebook_ neighbors are displayed in deep red and the most dis-similar <u>GI50</u>_codebook_ neighbors appear in bright yellow. Two distinctive features characterize SOM_DTP_. First, FDA approved compounds are projected on SOM_DTP_ as blue hexagons, where hexagons are sized according to the number of FDA agents appearing in any SOM_DTP_ node. Inspection of **Figure 2** finds a general tendency for approved agents to project to SOM_DTP_ nodes with unique patterns (e.g. dissimilar <u>GI50</u>_codebooks_). Statistical support for this observation is displayed in the right half of **Figure 2,** in the form of histograms based in inter-node <u>GI50</u>_codebook_ distances for nodes containing FDA approved agents (top right panel) and lacking FDA approved agents (bottom right panel). A pairwise t-test for the vector distance between these two groups finds a p-value of 5.3e-7, in support of the visual association of FDA compounds and unique (e.g dis-similar) chemosensitivity patterns. Second, inspection of compound names for FDA screened agents are displayed across the SOM_DTP_ in **Figure 2**. A listing of these nodes and their SOM_DTP_ projections appear in **Appendix Table III**. In summary, FDA compounds with known MOAs are grouped together, with, for example, nucleic acid targeting agents appearing in the upper right corner of SOM_DTP_, tubulin targeting agents at the left edge slightly below the horizontal dividing line and defective BRAF targeting agents appearing near the middle. Collectively, these results support our prior report [7] of associations between screened compounds, their MOAs and projections on SOM_DTP_.

While each SOM_DTP_ node represents a cluster of <u>GI50</u>_NCI60_ values, a more global, lower resolution representation, that optimally groups SOM_DTP_ nodes into meta-clades, is proposed. Three procedures were used for determining the optimal number of meta-clades; the **gap_statistic** method determines the total within intra-cluster variation for different numbers of meta-clades (selecting the cluster size that maximizes the gap_statistic), the **silhouette** method computes the average silhouette of observations for different numbers of meta-clades (selecting the optimal cluster size that maximizes the average silhouette) and the within-custer sum of squares method (**WSS**) minimizes the within-meta-clade sum of squares (a measure of within meta-clade similarity) and maximizes the between-meta-clade sum of squares (a measure of how separated each meta-clade is from the others). **Figure 3** displays the results for these three methods using <u>GI50</u>_codebooks_. The optimal cluster size is indicated by the vertical lines in each plot. The **silhouette** and **gap_statistic** methods yield 28 clusters as optimal, while the WSS method finds 26 as the optimal cluster size for this data. A value of 28 was selected for the optimal number of meta-clades used in this analysis.

**Figure 3.**
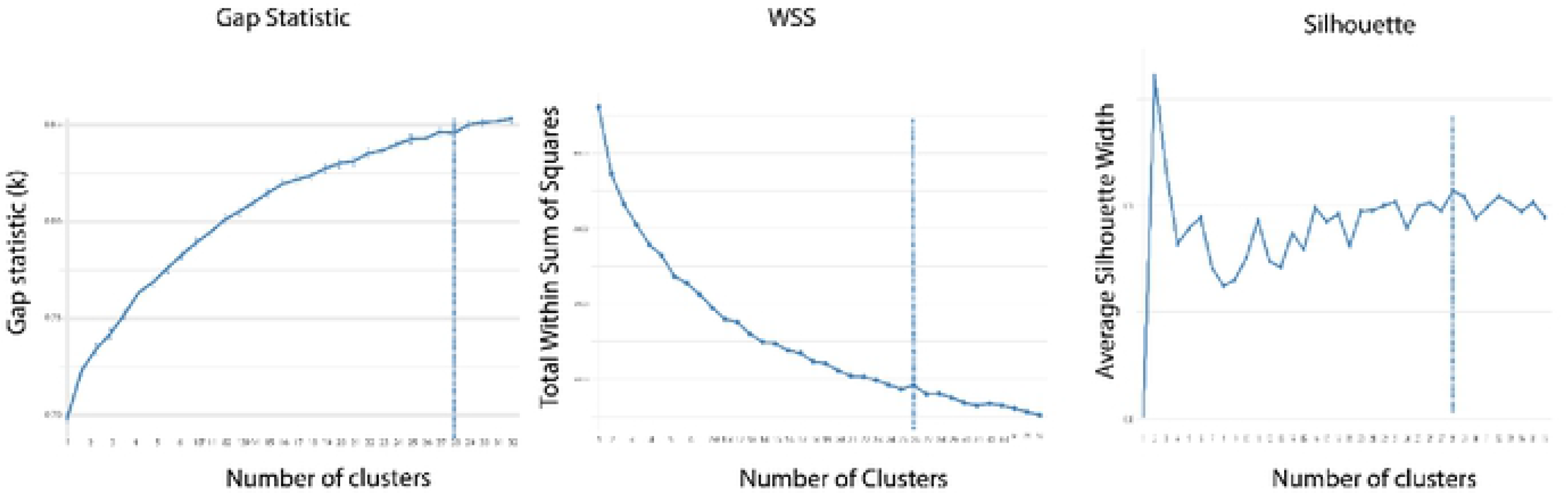
Results for selecting optimal cluster size using the gap_statistic (left panel), within sum of squares (middle panel) and silhouette (right panel) methods. The gap statistic and within sum of squares methods rely on an inflection point on each curve for optimal cluster sizewhile the silhouette methods seeks the largest value for silhouette width. These results indicate an optimal number of clusters in the 26-28 range.

A visual perspective these 28 meta-clades appears in **Figure 4**. The 1232 <u>GI50</u>_codebooks_ appear as a clustered heatmap (Euclidean,Ward’s) in the left panel of **Figure 4**. The left-most dendrogram, adjacent to this heatmap, displays the clade tree for hierarchical clustering of <u>GI50</u>_codebooks_. The cutree tool [15] is used to group the dendrogram into 28 meta-clades. The vertical ribbon, adjacent to this dendrogram, colored spectrally from blue to red, represents a subdivision of the hierarchal clade tree into 28 meta-clades. The pvclust utility [16], using random resampling, confirms this set of 28 meta-clades with a confidence p-value above 0.995 across resampling (n=1000 resamples). The right panel in **Figure 4** displays the 28 meta-clades on SOM_DTP_, colored according to the spectral-colored ribbon in the left panel. The data reduction of 1232 SOM_DTP_ nodes to 28 meta-clades yields a lower resolution, more manageable, perspective of the complete SOM_DTP_. SOM_DTP_ meta-clades will be assessed according to their chemosensitivity of tumor cells with and without defective genes. Rather than report results for all 28 meta-clades, results will be consolidated in to 7 meta-clade groups. The cutree utility, specifying 7 groupings of the complete dendrogram, will be used. Groupings consist of **A**: meta-clades 1-6, **B**: meta-clades 7-9, **C**: meta-clades 10-15, **D**: meta-clades 16-18, **E**: meta-clades 19-20, **F**: meta-clades 21-24 and **G**: meta-clades 25-28. The vertical grayscale colored bar in **Figure 4** displays from these groupings from **A** (bottom:black) to **G** (top:light gray).

**Figure 4:**
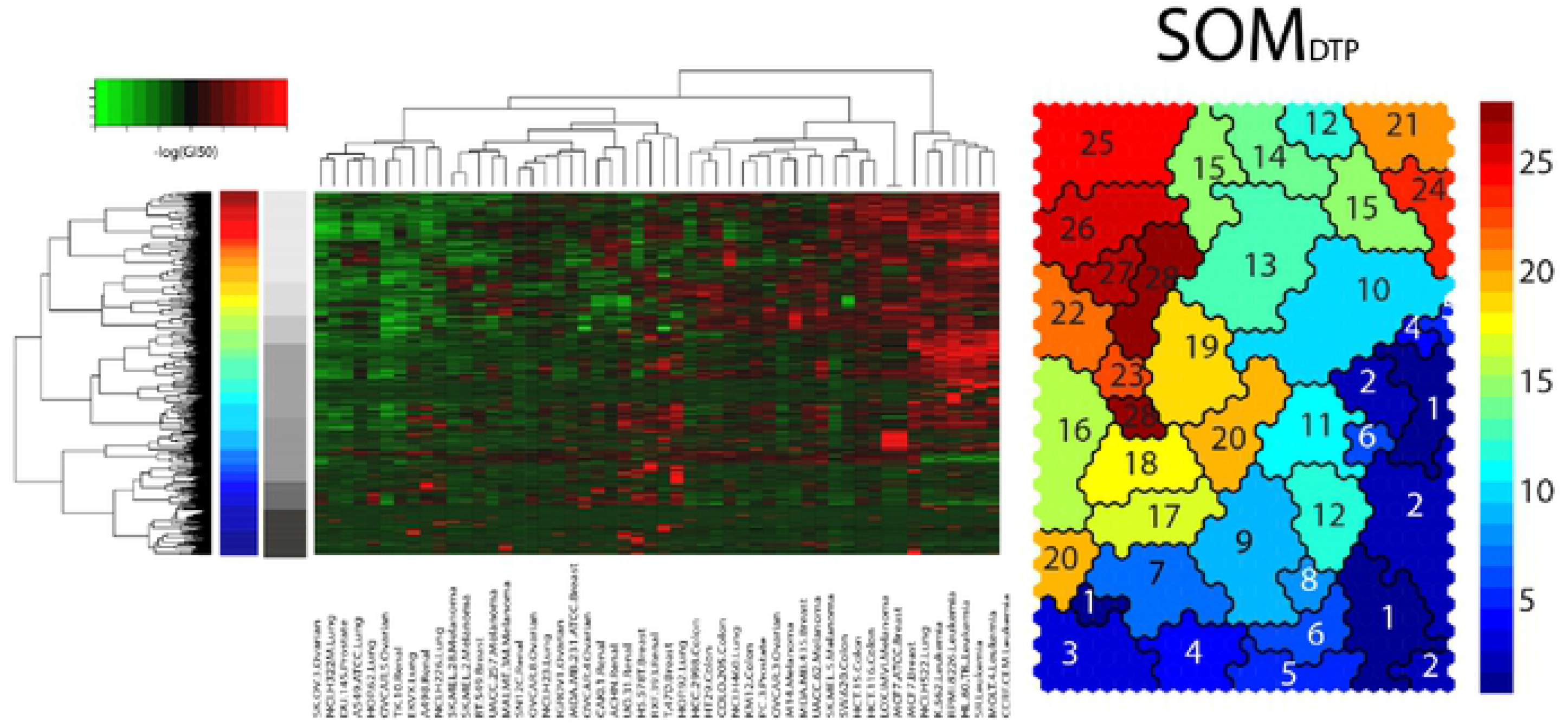
Left panel displays a heatmap of <u>GI50</u>_codebook_, colored spectrally from green(chemoinsensitive) to red(chemosensitive). Dendrogram at the left represents hierarchical clustering (Euclidean, Ward’s) of <u>GI50</u>_cooebooks_. Right panel displays the SOM_DTP_ colored according to hierarchical cutree (15) specified at the optimal number of 28 meta­ clades. The 28 colors appear spectrally from meta-clade 1 (dark blue), at the bottom of the hierarchical dendrogram, tometa-clade 28 (dark red), at the top of the hierarchical dendrogram. Grayscale bar adjacent to the 28 meta-clade spectrally colored bar displays the 7 meta clades groupings to be analyzed.

Noteworthy is the mapping of the 28 cutree clades to discontinuous SOM_DTP_ regions. Ideally, cutree clades would appear as contiguous regions the 2-dimensional SOM_DTP_. However, this is not the case. To obtain contiguous SOM_DTP_ regions, an alternative hierarchical clustering algorithm would need to be used that only combines adjacent dendrogram clades that appear beside each other on SOM_DTP_. Assigning contiguous SOM regions is an active area of research into dimensionality reduction [https://en.wikipedia.org/wiki/Nonlinear_dimensionality_reduction], with specific focus on representing SOMs in one dimension https://www2.cs.arizona.edu/~paolosimonetto/Papers/EuroVis14_1_slides.pdf [17]. Many of these efforts use randomized resampling to identify contiguous map regions by consensus. Usually standard hierarchical clustering suffices, and any outlying (noncontiguous) points can be accounted for manually. Towards this end, SOM_DTP_ singletons, appearing as a hierarchical clade that maps to SOM_DTP_ as a node without the same meta-clade neighbors, have been replaced by their neighborhood meta-clade assignments. There are 12 such cases for this data set. An additional explanation for non-contiguous SOM_DTP_ meta-clades involves differences in methodology. SOMs organize data by mapping each cluster to its most similar neighbors (six in the case of hexagonal mapping); whereas hierarchical clustering, as used to obtain this heatmap, builds each branch of the dendrogram by pairwise associations. A further additional consideration is due to assignments of distances used for clustering (Euclidean for hierarchical clustering and Epanechnikov Function[8] for SOMs). Our choice of the Epanechnikov Function for SOM clustering consistently yielded the lowest SOM_DTP_ quantization errors [7].

Linking defective genes to <u>GI50</u>_NCI60_ begins by partitioning each <u>GI50</u>_codebook_ into subsets comprising tumor cells with a defective gene or gene combination (e.g. <u>GI50</u>_defective_) and those lacking these alterations (<u>GI50</u>_wt_) and applying a Students t-test to identify cases of relative chemosensitivity (**Box D** in **Figure 1**). An example illustrates this process. **Figure 5** summarizes results for the defective ABL1 gene tested for <u>GI50</u>_codebook_ at SOM_1,13_ (subscripting refers to the SOM node, i.e. SOM_row,column_). Five NCI60 tumor cells harbor this altered gene, with these tumor cells having a mean <u>GI50</u>_codebook_ response of 0.97 compared to −0.11 for the wild type tumor cells (p=6.91e-3). The upper panel in **Figure 5** displays <u>GI50</u>_codebook_, ordered from most chemosensitive to least chemosensitive values. NCI60 tumor cells with the ABL1 alteration, highlighted in red and representing <u>GI50</u>_defective_, are ranked at positions 3, 6, 8, 25 and 51. The lower portion of **Figure 5** displays the <u>GI50</u>_component_ for each of the 5 tumor cells having the ABL1 gene alteration. Recall that the 1232 <u>GI50</u>_component_ values can be displayed in the 44X28 SOM_DTP_ format. Regions of greatest and least chemosensitivity are displayed spectrally from red to blue. Noteworthy is the projection of Gleevec chemosensitivity to the most sensitive <u>GI50</u>_component_ regions (e.g.red) for K-562, RMPI-8226 and HS578T.

**Figure 5.**
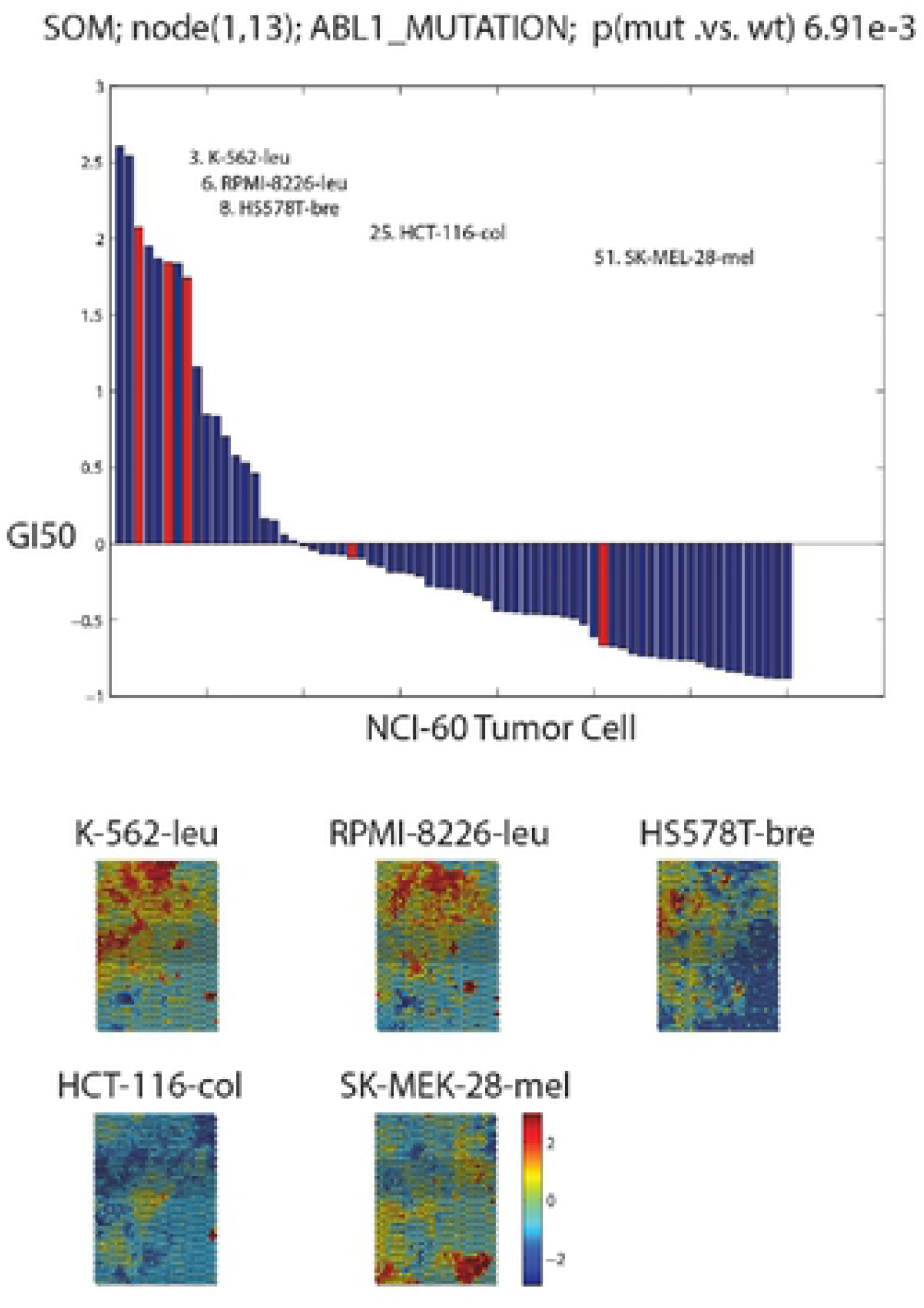
Upper panel displays <u>GI50</u>_codebook_ for SOM_1, 13_, ordered from most to least chemosensitivity. The 5 tumor cells with the defective ABLl gene appear as red bars. Lower panel displays <u>GI50</u>_component_ for the 5 tumor cells with defective ABLl. SOM nodes are colored spectrally from highest chemosensitivity (red) to lowest chemosensitivity (blue).

A Students t-statistic represents the significance comparing the chemosensitivity of a SOM node for tumor cells with, versus without, defective genes for each SOM_DTP_ node (SOM_1,13_ in the above example for ABL1). The left panel in **Figure 6** displays the Students t-statistic for defective ABL1 on SOM_DTP_; where t-statistic values are colored spectrally from low(blue) to high(red) significance. SOM_DTP_ nodes without statistical significance are not colored. The most significant t-statistics for defective ABL1 are located mainly in SOM meta-clades 1, 10, 14 and 26. Gleevec appears as the most significant SOM_5,15_ node in meta-clade 14. For comparison, the results for KRAS are displayed in the middle panel of **Figure 6**. Here there are 12 tumor cells (A549/ATCC-Lung, CCRF-CEM-Leukemia, HCC-2998-Colon, HCT-116-Colon, HCT-15-Colon, HOP-62-Lung, NCI-H23-Lung, NCI-H460-Lung, OVCAR-5-Ovarian, RPMI-8226-Leukemia, SK-OV-3-Ovarian and SW-620-Colon) harboring defective KRAS, with the significant chemosensitive SOM_DTP_ nodes appearing in meta-clades 21, 22 and 27. SOM meta-clade 21 is the location of cytarabine (ara-C) and is consistent with the conclusion of Ahmad et al [18] that adult AML patients carrying defective KRAS benefit from higher ara-C doses more than wt KRAS patients.

**Figure 6:**
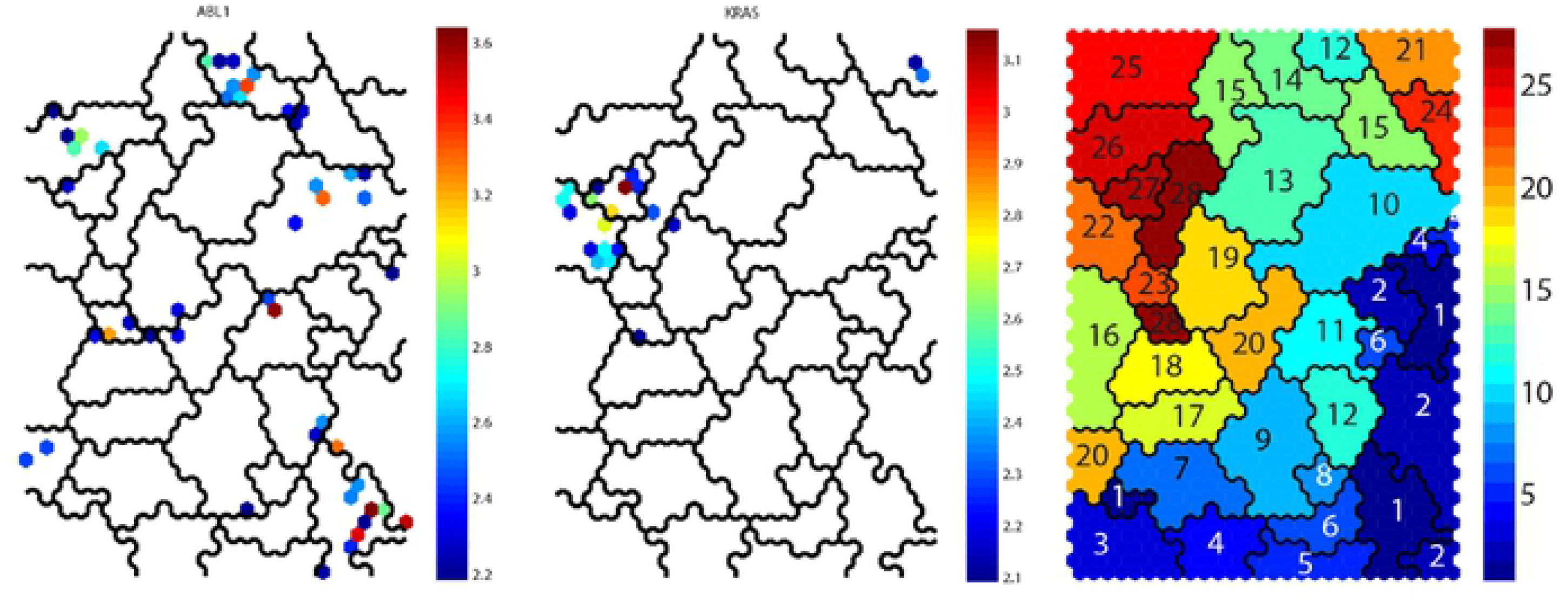
Left and middle panels display significant chemosensitive (blue:least, red:most) SOM_DTP_ nodes for tumor cells with defective ABLl and KRAS, respectively. Rightpanel displays SOM meta-clades.

**Appendix Figures 3** and **4** display additional examples for the defective genes PIK3RI and IGF1R, respectively. PIK3R1 (Phosphatidylinositol 3-Kinase Regulatory Subunit Alpha) and a related gene, PIK3CA (PI3-Kinase Subunit Alpha) are lipid kinases capable of phosphorylating the 3’OH of the inositol ring of phosphoinositides. Both are responsible for coordinating a diverse range of cell functions including proliferation and survival. Defective PIK3CA has been documented by Holbeck et al.[19] to enhance tamoxifen sensitivity in selected NCI60 tumor cells. The results here also find chemosensitivity in NCI60 tumor cells having defective PIK3R1. The second example of defective IGF1R supports the importance of evaluating drug sensitivity for compounds targeting leukemia cells [20] and the emergence of IGF1R as a potential therapeutic target for the treatment of different types of cancer including plasma cell myeloma, leukemia, and lymphoma [21]. Both examples illustrate potential roles of defective genes in chemosensitivity.

SOM_DTP_ projections for the most frequent primary CellMiner MOA assignments (Tu, T1, T2, A, D, Apo and PK) are displayed in the left panel of **Figure 7**. Inspection indicates that MOA classes A, D, T1 and T2 appear mainly in the upper right SOM region (SOM meta-clade 21; Group **A**), while MOA Apo appears mainly in the upper left region (SOM meta-clades 25 and 26; Group **G**). Tu compounds are found mainly in SOM meta-clades 16, 17 and 18 (Group **D**). SOM meta-clade 19(Group **E**) consists of only MOA PK; while MOA PK compounds are in the majority for SOM meta-clades 1 through 6(Group **A**). The horizontal gray scale bar at the bottom of the right panel in **Figure 4** identifies the seven meta-clade groups assigned earlier. Inspection indicates relative similarities of MOA types within each of the seven meta-clade groups **A**:**G**. Notable is the majority representation of MOA:PK in Group **A** and MOA:Tu in Group **D**. Detailed results for MOAs within meta-clade groups will be presented below.

**Figure 7:**
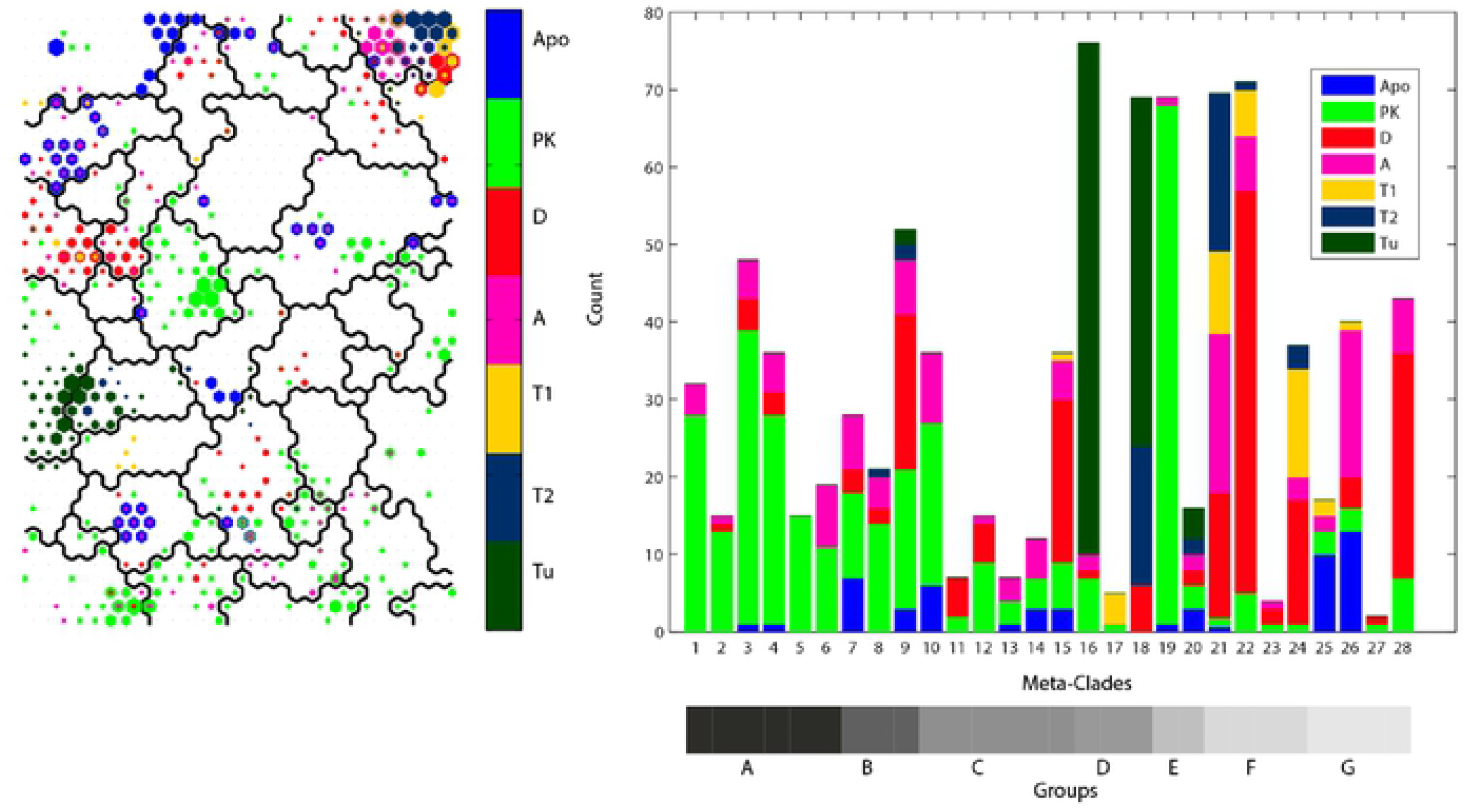
Left panel; SOM_DTP_ projections for FDA approved compounds for the primary CellMiner assigned MOAs. Right panel display a histogram for counts of MOA classes within SOM_DTP_ meta-clades. Projections include the top 8 SOM_DTP_ nodes for each compound. Horizontal grayscale bar below the right-hand panel indicates meta-clade groups A:G.

The right panel in **Figure 7** displays a histogram of the counts for these primary MOAs across SOM meta-clades. Primary MOAs appear color-coded in each vertical bar, with their heights corresponding to MOA counts in each meta-clade. **Table I** lists the most frequent primary MOAs for meta-clade groups A:G. These results segregate the primary MOAs into separate regions of the hierarchical dendrogram; MOA:PK appears at the bottom, MOAs targeting DNA and Apo appear at the top and mixtures of primary MOAs appear in the middle. These meta-clade grouping will be analyzed in greater detail for links of MOAs to defective genes.

**Table I.** Most frequent primary MOA assignments within meta-clade groups **A**:**G**

| Meta-clade Groups | MOA |
| --- | --- |
| A: 1-6 | PK |
| B: 7-9 | PK, A |
| C: 10-15 | PK, A, D |
| D: 16-18 | Tu, T2 |
| E: 19-20 | PK |
| F: 21-24 | A, D, T1 |
| G: 25-28 | Apo, A, D |

Fisher’s exact scores are used to identify cases where the SOM_DTP_ projections of defective genes are statistically enriched in co-projections of MOA types. **Figure 8** displays a bar chart of the log(pvalue) for significant (p<0.05) Fisher’s exact scores based on analysis of the complete SOM_DTP_, regardless of meta-clade identity. Forty-eight defective genes have significant Fisher’s exact scores in SOM_DTP_. Defective genes with the top-most significance scores include **AXL**, **STAT2**, **TNKS**, **MYC**, **ZHX2**, **CDKN2A**, **SPTBN1**, **NOTCH1**, **EGFR**, **RPTOR**, **MECOM** and **BRAF**. A Fisher’s exact test, restricted to each of the 28 meta-clades, will be used to identify defective genes within meta-clades.

**Figure 8:**
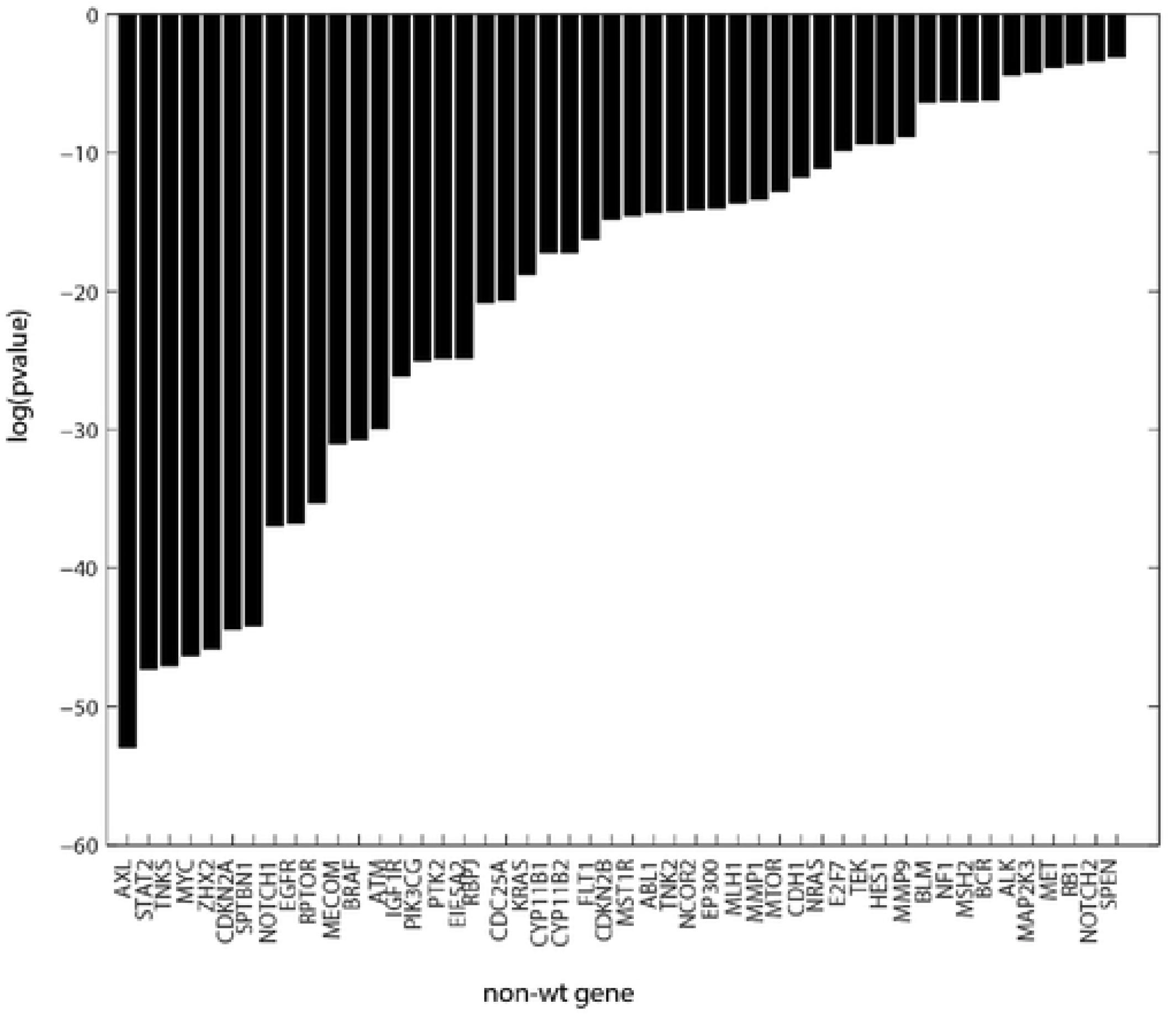
Fisher’s exact scores (log(pvalue), pvalue<0.05). Results are based on classifications using up to the 8^th^ best SOM projection nodes. Forty eight defective genes have significant Fisher’s exact scores. The results provided below will focus on these defective genes.

### <u>RESULTS</u>: Group A (meta-clades 1 through 6)

The results for SOM meta-clade group **A** find nine defective genes with significant Fisher’s exact scores (**ABL1, CDC25A, CDH1, CDKN2A, IGF1R, MMP1, NCOR2, NF1** and **RPTOR**) associated with ten MOA classes (AM, Ang, BCR-ABL, Db, Hyp, Ho, NonCan, PARP, PK and Pase). These results are summarized in **Figure 9**. MOA:PK represents the highest count with **CDKN2A**, **MMP1, NF1** and **ABL1** as the next most frequent defective genes. The second highest count is for MOA:BCR-ABL, which is associated with defective genes **NF1** and **CDKN2A**. MOA:Ho is the next highest count, with defective genes **NF1**, **MMP1**, **NCOR2** and **ABL1**.

**Figure 9:**
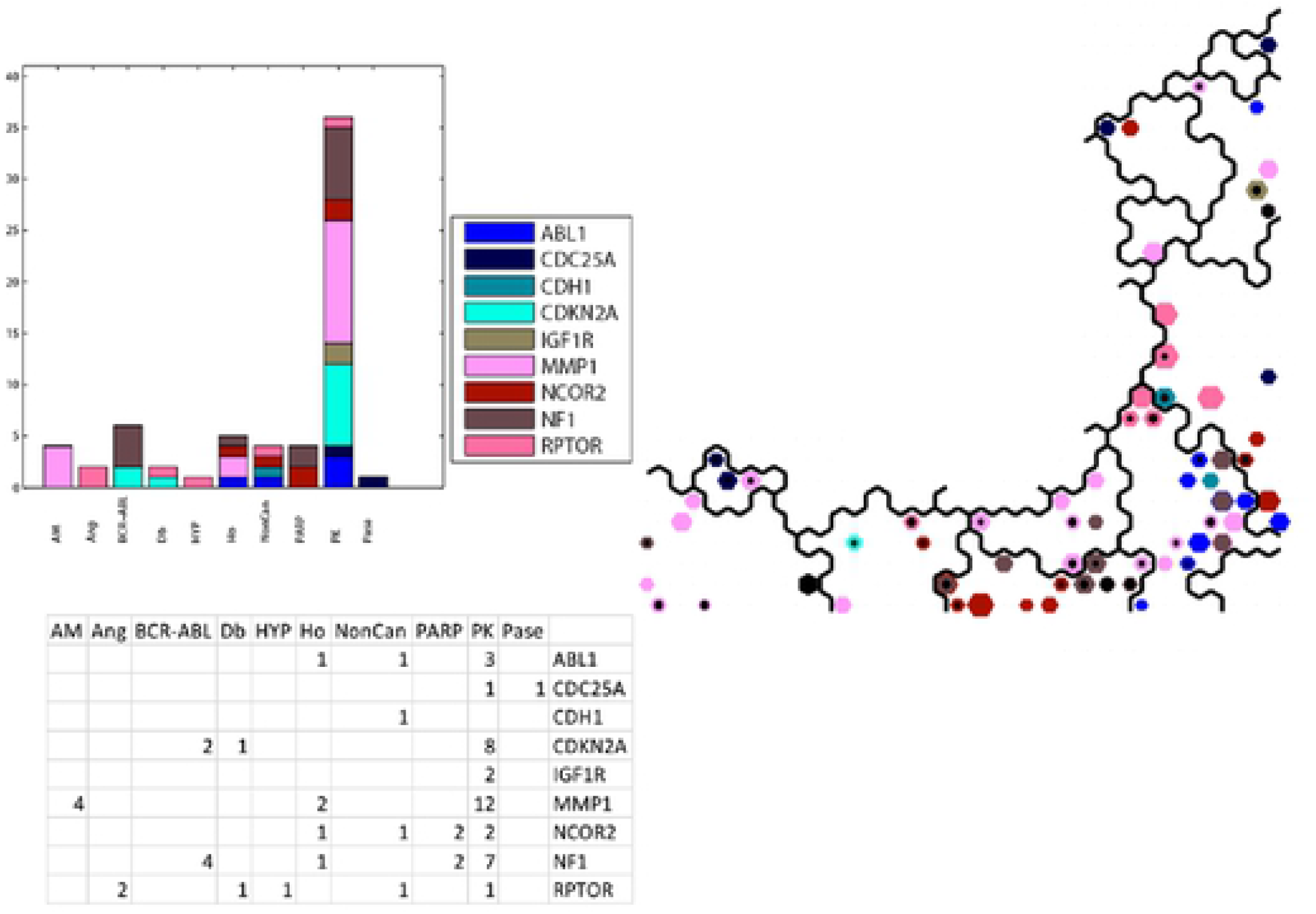
Upper left panel displays thehistogram of counts for the co-occurrence of SOM_DTP_ nodes with defective genes and FDA compound projections Lower left panel displays the counts for co occurrence. Right panel displays the SOM_DTP_ for meta cl ades 1 through 6 (Group A). SOM_DTP_ nodes displayed represent instances of nodes with significant t statistics when comparing SOM_codebooks_ grouped by tumor cells with and without defective genes. Node colors for defective genes correspond to the legend inserted into the upper left panel. Only co-occurrences for SOM_DTP_ projections of FDA compounds are displayed. The counts displayed in the lower left panel represent thetop-best 8 SOM_DTP_ projections of FDA compounds. A consistent coloring scheme is used for this and all subsequent figures, such that all defective genes presented in the **RESULTS** are assigned a unique color. Fisher’s exact statistics will use the top best eight SOM_DTP_ projections for FDA approved compounds.

The FDA compounds associated with the most frequent defective genes (**ABL1, NF1, CDKN2A and MMP1**) are summarized in **Table II**. All compounds within the MOA:PK class are associated either with **MMP1** or **NF1**. Only staurosporine (PK:PRKCA) is associated with all four genes, while MOA:Ho, MOA:NonCan and MOA:PARP represent less frequent cases.

**Table II.**
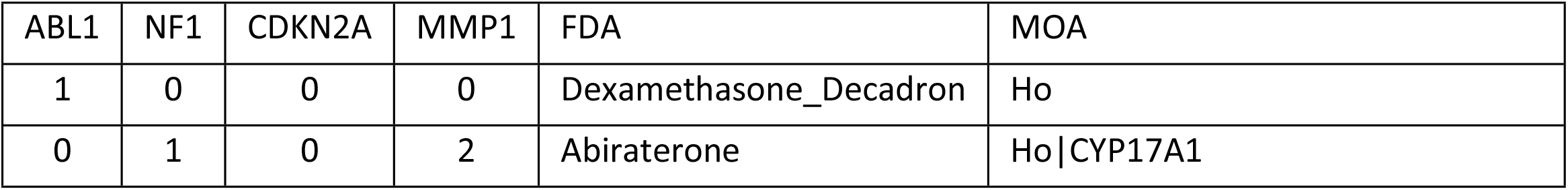
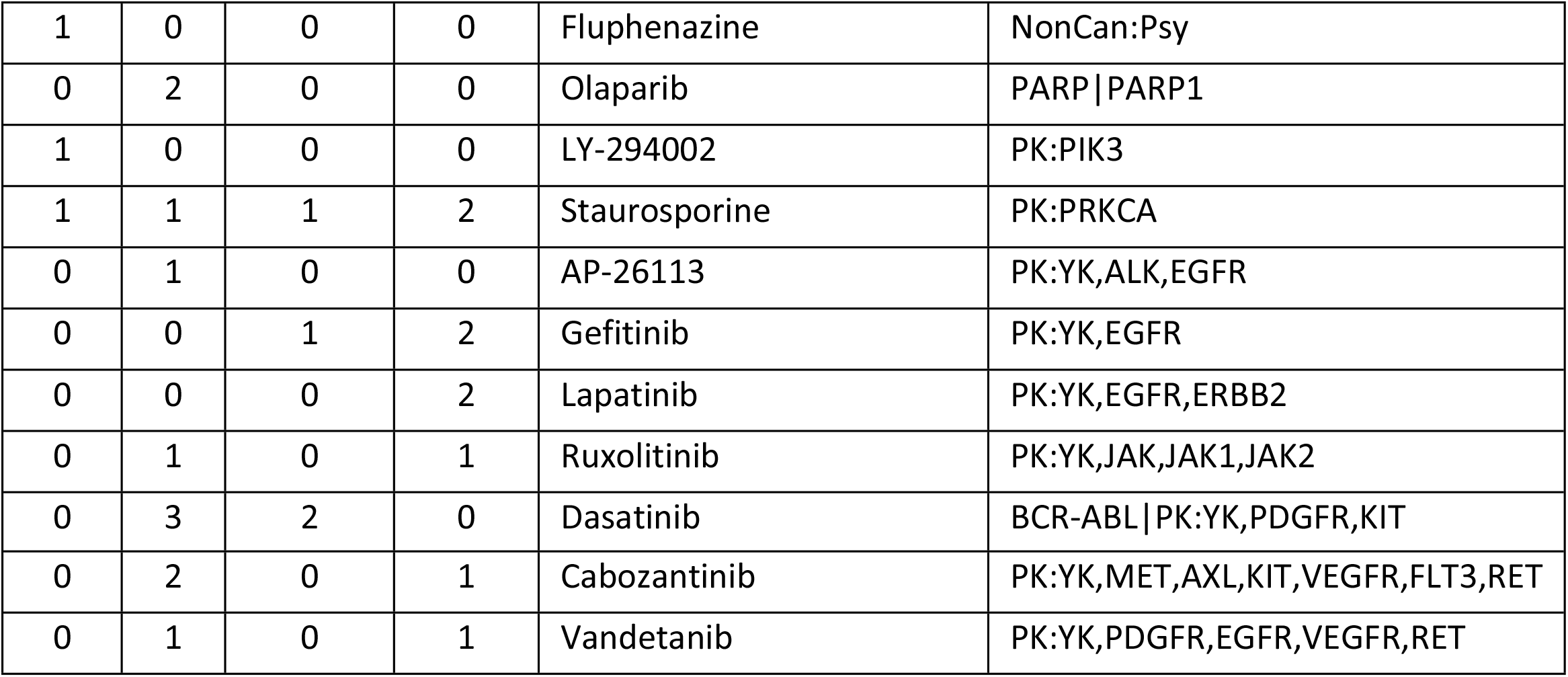
Group A(meta-clades 1 through 6). Table lists the defective genes that satisfy Fisher’s exact significance (p<0.05) for each meta-clade in Group A. Row entries list the counts for each FDA and MOA entry for significant defective genes.

| ABL1 | NF1 | CDKN2A | MMP1 | FDA | MOA |
| --- | --- | --- | --- | --- | --- |
| 1 | 0 | 0 | 0 | Dexamethasone_Decadron | Ho |
| 0 | 1 | 0 | 2 | Abiraterone | Ho\|CYP17A1 |
| 1 | 0 | 0 | 0 | Fluphenazine | NonCan:Psy |
| 0 | 2 | 0 | 0 | Olaparib | PARP\|PARP1 |
| 1 | 0 | 0 | 0 | LY-294002 | PK:PIK3 |
| 1 | 1 | 1 | 2 | Staurosporine | PK:PRKCA |
| 0 | 1 | 0 | 0 | AP-26113 | PK:YK,ALK,EGFR |
| 0 | 0 | 1 | 2 | Gefitinib | PK:YK,EGFR |
| 0 | 0 | 0 | 2 | Lapatinib | PK:YK,EGFR,ERBB2 |
| 0 | 1 | 0 | 1 | Ruxolitinib | PK:YK,JAK,JAK1,JAK2 |
| 0 | 3 | 2 | 0 | Dasatinib | BCR-ABL\|PK:YK,PDGFR,KIT |
| 0 | 2 | 0 | 1 | Cabozantinib | PK:YK,MET,AXL,KIT,VEGFR,FLT3,RET |
| 0 | 1 | 0 | 1 | Vandetanib | PK:YK,PDGFR,EGFR,VEGFR,RET |

The salient feature of the defective genes associated with SOM meta-clade Group **A** is their potential to influence the RAF/MEK/ERK and the PI3K/AKT pathways. For example, loss of **NF1** gene expression leads to increased RAS activation and hyperactivation of the downstream RAS effectors, including the RAF/MEK/ERK and the PI3K/AKT pathways [22]. Abnormal activation of RAS by defective **NF1** is a central driver event in some soft-tissue sarcomas (MPNST). Receptor tyrosine kinases (RTKs), including PDGFRA and EGFR, can activate RAS signaling and downstream factors such as MEK and mTOR. Ki et al. [23] find the addition of mTOR inhibitors to cells harboring defective **NF1** enhance the activity of DNA targeting agents. Defective genes that impact PI3K-Akt-mTOR signaling could weaken the tumor cell and enhance susceptibility to chemotherapeutic drugs.

A noteworthy entry in **Table II** is for Olaparib, MOA:PARP and defective gene **NF1**. Combination treatment with olaparib and various inhibitors of PD-L1, VEGFR, PI3K, and AKT may effectively inhibit the growth of rapidly proliferating TNBC cells[24]. A review of candidate synthetic lethality partners to PARP inhibitors in the treatment of ovarian clear cell cancer by Kawahara et al. [25] finds **PARP** and **NF1** to be synthetic lethality pairs [26]. Synthetic lethality (SL) describes the genetic interaction by which the combination of two separately non-lethal mutations results in lethality [27]. Generally, the ablation of two genes located in parallel pathways (leading to cell survival or a common essential product) is one of the important patterns causing synthetic lethality. Synthetic lethality appears to be achieved with combined EGFR and PARP inhibition [28]. SL has recently emerged as a promising new approach to cancer therapy [29].

A direct role of **MMP1** on chemosensitivity in SOM meta-clade Group **A** has not been reported. However, Zhou et al. [30] identify **MMP1** as a potential gene conferring resistance of EGFR drugs targeting in non-small cell lung cancer. Rapamycin significantly enhanced the expression of interstitial collagenase (**MMP1**) at the protein and mRNA levels [31]. An assessment of upregulated expression levels in serous ovarian cancer cells by Zhang et al. [32] find matrix metalloproteinase 1 (**MMP1**) to be among the most upregulated mRNAs in the chemoresistant cell lines. Given that **MMP1** is the most frequent defective gene associated with MOA:PK (cf. **Figure 9**), combined with its role in chemosensitivity, suggests that defective **MMP1** may play a role in the weak <u>GI50</u>_NCI60_ responses to PIK3 and EGFR agents screened in the NCI60.

Defective **CDKN2A** represents the second most frequent defective gene within MOA:PK and MOA:BCR-ABL. The importance of defective **CDKN2A** will be provided in the analysis of meta-clade group **F**, where it has the highest frequency of counts for numerous MOA classes.

### <u>RESULTS</u>: Group B (meta-clades 7 through 9)

Eleven MOA classes (A7, AM, Apo, Db, Ds, HDAC, Ho, NonCan, PARP, PK and T) are associated with fifteen defective genes (**AXL, E2F7, EIF5A2, EP300, HES1, IGF1R, MECOM, MMP1, MYC, NRAS, PIK3CG, PTK2, TNK2, TNKS** and **ZHX2**) for SOM meta-clades 7 through 9. **Figure 10** summarizes these results.

**Figure 10.**
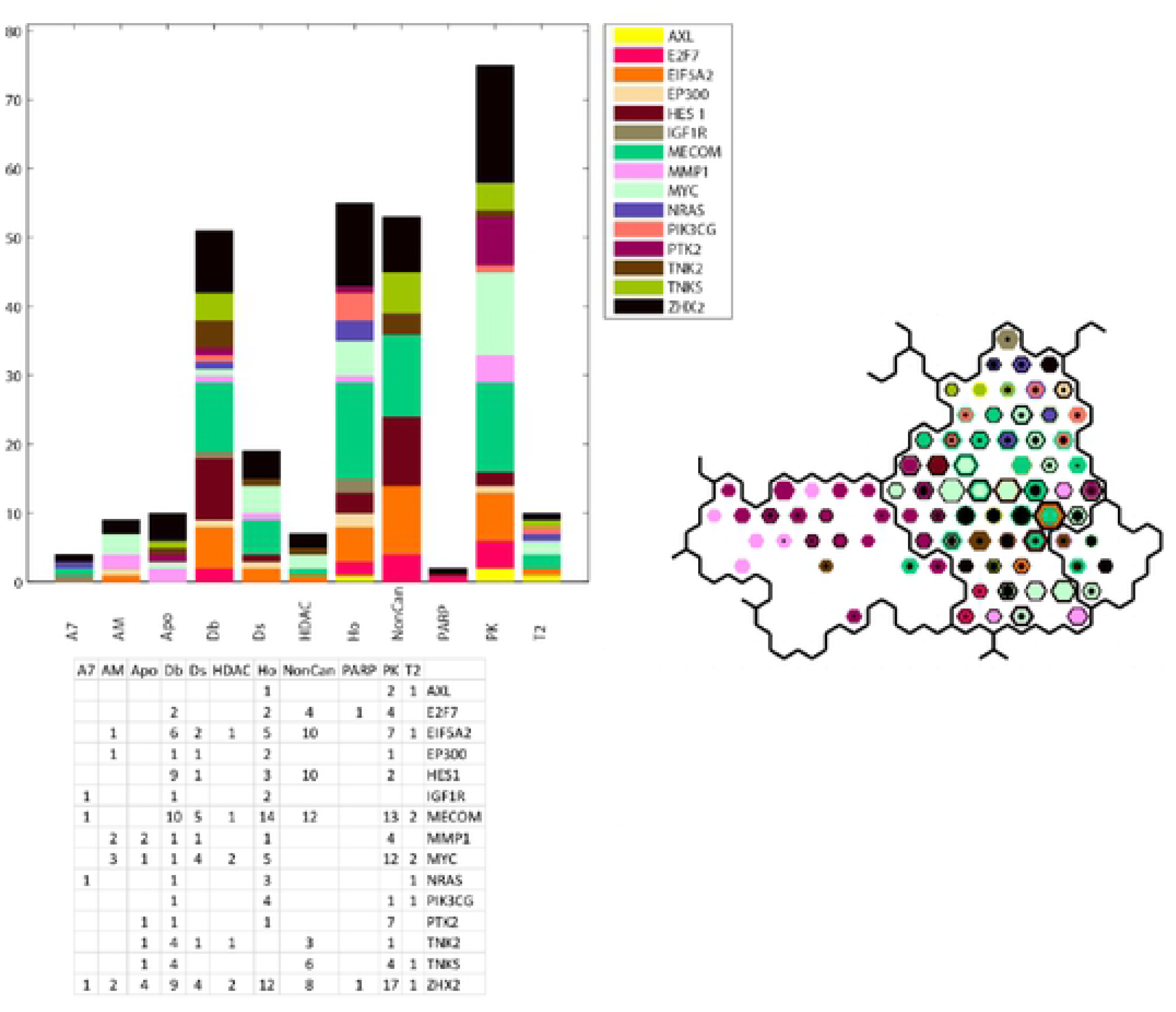
Upper left panel displays the histogram of counts for the co-occurrence of SOM_DTP_ nodes with defective genes and FDA compound projections Lower left panel displays the counts for co occurrence. Right panel displays the SOM_DTP_ for meta cl ades 7 through 9 (Group **B**). SOM_DTP_ nodes displayed represent instances of nodes with significant t-statistics when comparing <u>GI50</u>_cooebooks_ grouped by tumor cells with and without defective genes. Node colors for defective genes correspond to the legend inserted into the upper left panel. Only co occurrences for SOM_DTP_ projections of FDA compounds are displayed. The counts displayed in the lower left panel represent the top-best 8 SOM_DTP_ projections of FDA componds.

The most frequent of these fifteen defective genes with significant Fisher’s exact scores are **MECOM, MYC, EIF5A2, HES1**, **TNKS**, **E2F7** and **PTK2** (**Table III).** The salient feature of these five defective genes associated with SOM meta-clades 7 through 9 is their potential to influence cellular functions such as protein synthesis, differentiation and translation. Recently deregulation of the protein synthesis apparatus has begun to gain attention as a major player in cancer development and progression. Among the numerous steps of protein synthesis, deregulation of the process of translation initiation appears to play a key role in cancer growth and proliferation [33]. Eukaryotic translation initiation factor (**EIF5A**) stimulates protein synthesis [34]. The interplay between **E2F7** and ribosomal rRNA gene transcription regulates protein synthesis [35]. Eukaryotic translation initiation factor 5A-2 (**EIF5A**) is involved in doxorubicin-induced epithelial-mesenchymal transition of cancer cells [36]. E2F transcription factors, such as **E2F7**, play essential roles in the regulation of cell cycle progression [37]. **HES1** is one mammalian counterpart of the Hairy and Enhancer of split proteins that play a critical role in many physiological processes including cellular differentiation, cell cycle arrest, apoptosis and self-renewal ability[38]. Up-regulation of **HES1** involves canonical or non-canonical pathways, such as the Hedgehog, Wnt and hypoxia pathways, frequently aberrant in cancer cells, found to be important in the maintenance of epithelial–mesenchymal transition (EMT) process induction. **MECOM** is found to be commonly enriched in cancer cells. Makondi et al. [39] suggest that targeting the MAPK signal transduction pathway through the targeting of the **MECOM** might increase tumor responsiveness to irinotecan treatment. **TNKS** promotes aerobic glycolysis and proliferation of ovarian cancer through activation of Wnt/β-catenin signaling [40]. **TNKS** inhibitor XAV939 increases chemosensitivity in colon cancer cell lines via inhibition of the Wnt signaling pathway [41]. A therapeutic strategy proposed by Thorvaldsen et al., targets TNKS to treat Wnt-dependent tumors [42]. **PTK2** (FAK), a protein tyrosine kinase in the RTK pathway, is an important mediator within the cell migration process, as well as in cell motility, survival and proliferation through kinase-dependent and -independent mechanisms [43]. Because of the involvement of **PTK2** (FAK) in many cancers, drugs that inhibit FAK are being sought and evaluated[44]. A screen to identify mechanisms of bleomycin resistance identified Sky1, **PTK2** and Agp2 as determinants of chemosensitivity[45].

**Table III.**
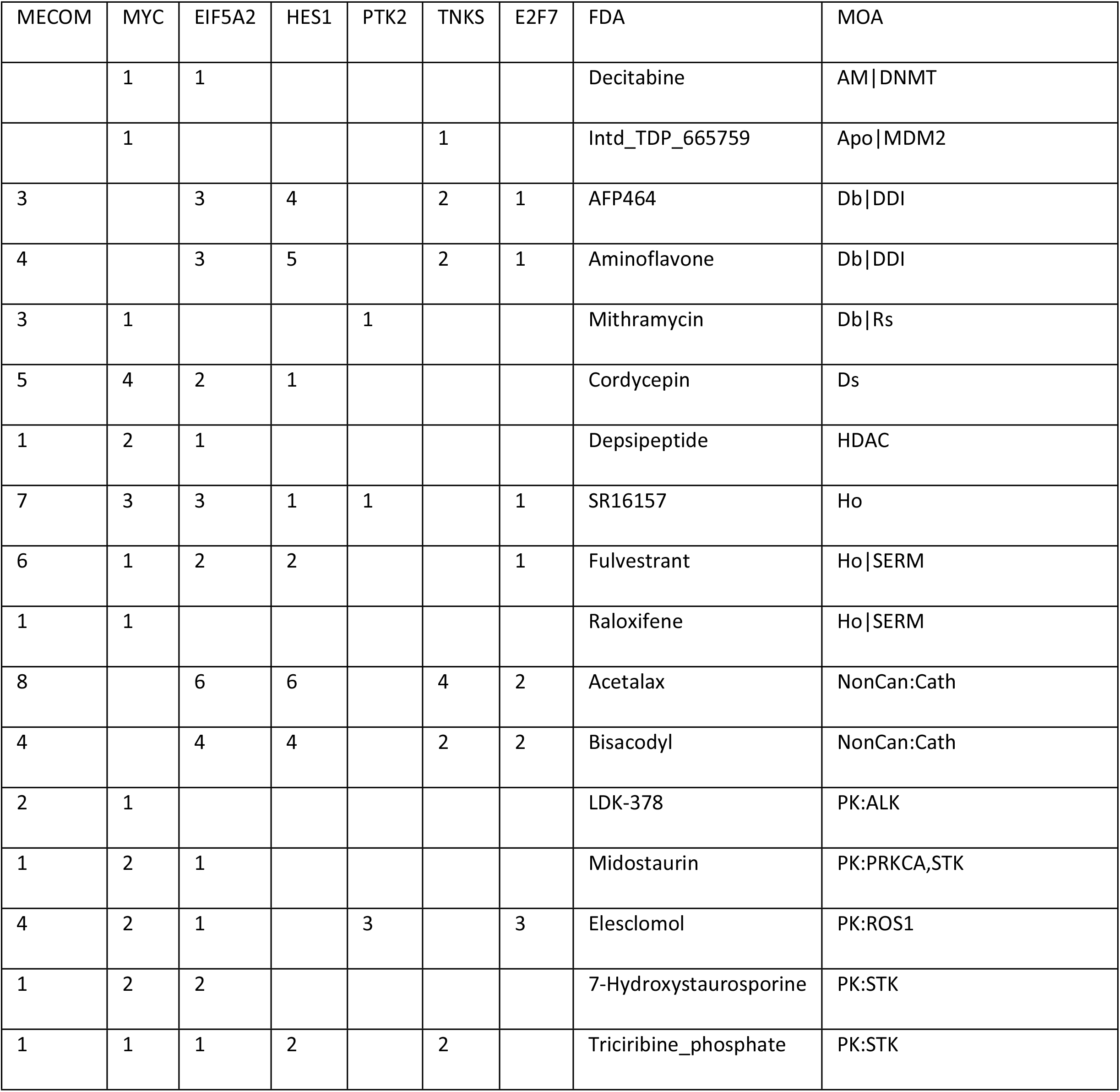
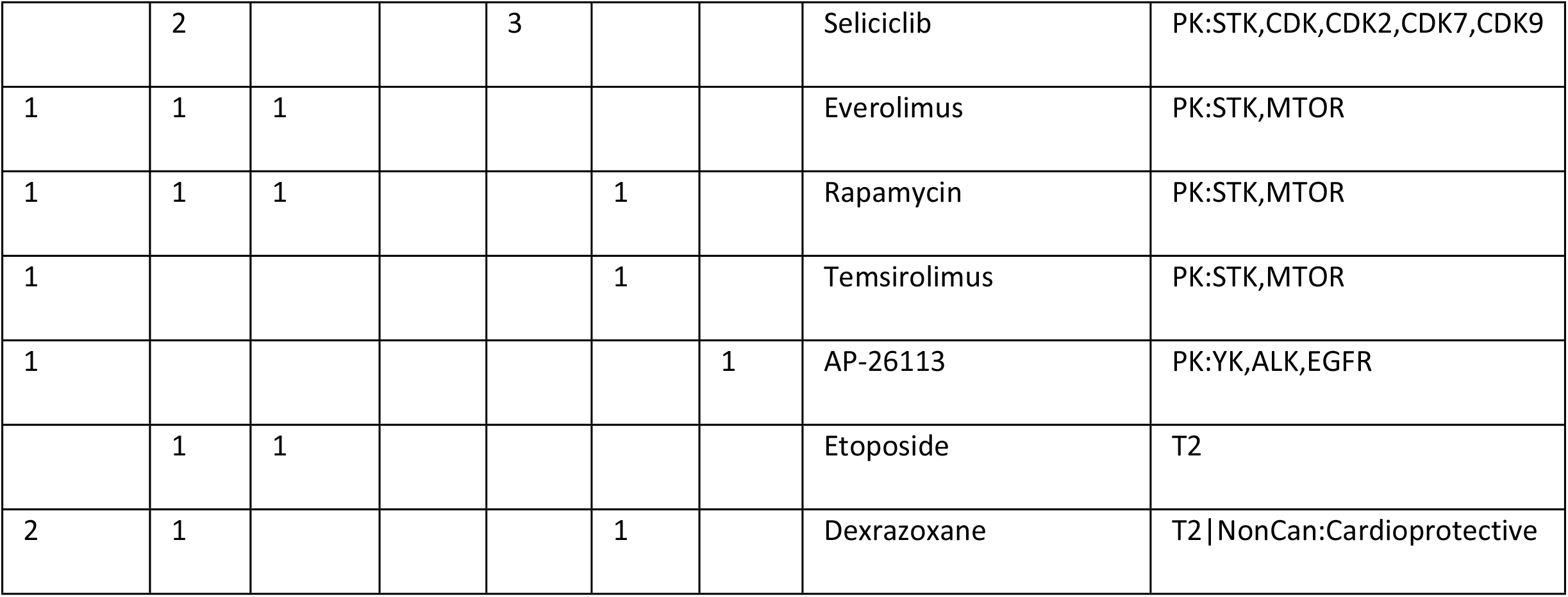
Group **B** (meta-clades 7 through 9). Table lists the defective genes that satisfy Fisher’s exact significance (p<0.05) for each meta-clade in Group **B**. Row entries list the counts for each FDA and MOA entry for significant defective genes.

| MECOM | MYC | EIF5A2 | HES1 | PTK2 | TNKS | E2F7 | FDA | MOA |
| --- | --- | --- | --- | --- | --- | --- | --- | --- |
|  | 1 | 1 |  |  |  |  | Decitabine | AM\|DNMT |
|  | 1 |  |  |  | 1 |  | Intd_TDP_665759 | Apo\|MDM2 |
| 3 |  | 3 | 4 |  | 2 | 1 | AFP464 | Db\|DDI |
| 4 |  | 3 | 5 |  | 2 | 1 | Aminoflavone | Db\|DDI |
| 3 | 1 |  |  | 1 |  |  | Mithramycin | Db\|Rs |
| 5 | 4 | 2 | 1 |  |  |  | Cordycepin | Ds |
| 1 | 2 | 1 |  |  |  |  | Depsipeptide | HDAC |
| 7 | 3 | 3 | 1 | 1 |  | 1 | SR16157 | Ho |
| 6 | 1 | 2 | 2 |  |  | 1 | Fulvestrant | Ho\|SERM |
| 1 | 1 |  |  |  |  |  | Raloxifene | Ho\|SERM |
| 8 |  | 6 | 6 |  | 4 | 2 | Acetalax | NonCan:Cath |
| 4 |  | 4 | 4 |  | 2 | 2 | Bisacodyl | NonCan:Cath |
| 2 | 1 |  |  |  |  |  | LDK-378 | PK:ALK |
| 1 | 2 | 1 |  |  |  |  | Midostaurin | PK:PRKCA,STK |
| 4 | 2 | 1 |  | 3 |  | 3 | Elesclomol | PK:ROS1 |
| 1 | 2 | 2 |  |  |  |  | 7-Hydroxystaurosporine | PK:STK |
| 1 | 1 | 1 | 2 |  | 2 |  | Triciribine_phosphate | PK:STK |
|  | 2 |  |  | 3 |  |  | Seliciclib | PK:STK,CDK,CDK2,CDK7,CDK9 |
| 1 | 1 | 1 |  |  |  |  | Everolimus | PK:STK,MTOR |
| 1 | 1 | 1 |  |  | 1 |  | Rapamycin | PK:STK,MTOR |
| 1 |  |  |  |  | 1 |  | Temsirolimus | PK:STK,MTOR |
| 1 |  |  |  |  |  | 1 | AP-26113 | PK:YK,ALK,EGFR |
|  | 1 | 1 |  |  |  |  | Etoposide | T2 |
| 2 | 1 |  |  |  | 1 |  | Dexrazoxane | T2 \| NonCan:Cardioprotective |

### <u>RESULTS</u> Group C (meta-clades 10 through 15)

Fourteen MOA classes (A7, AM, Apo, BCR_ABL, BRD, Ds, HDAC, Ho, Mito, NFKB, PARP, PK, PSM and T1) are associated with eleven defective genes (**ATM, BRAF, CDC25A, EGFR, IGF1R, MLH1, NCOR2, NOTCH1, PIK3CG, SPTBN1 and STAT2**) for meta-clades 10 through 15 that have a significant Fisher’s exact score. **Figure 11** summarizes these results.

**Figure 11.**
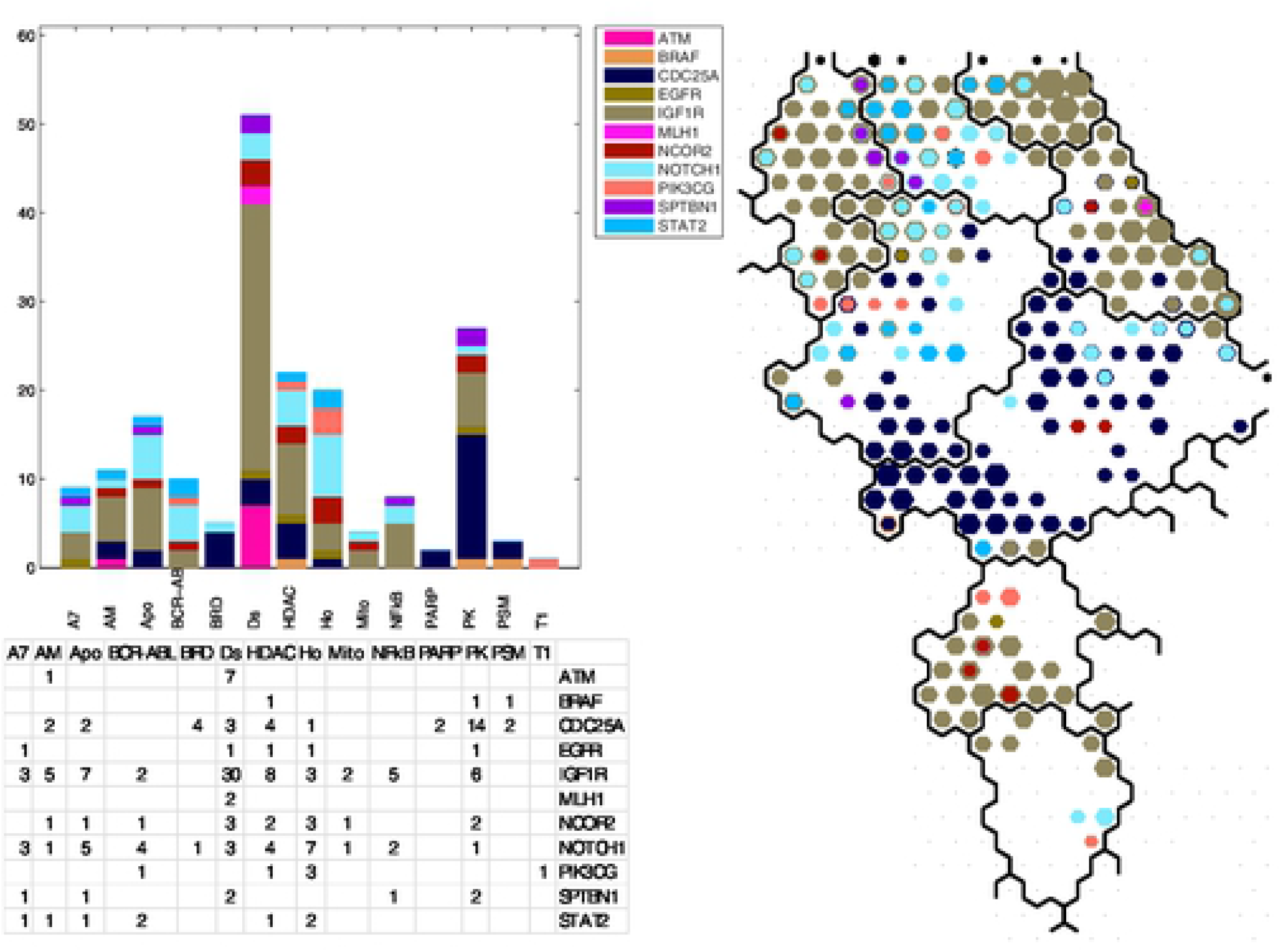
Upper left panel displays the histogram of counts for the co occurrence of SOM_DTP_ nodes with defective genes and FDA compound projections. Lower left panel displays the counts for co occurrence. Right panel displays the SOM_DTP_ for meta clades 10through 15 (Group C). See legend to **Figure 9** for additional details.

The most frequent defective genes in meta-clades 10 through 15 are **IGFR1, ATM, NOTCH1, CDC25A and BRAF** (**Table IV).** The salient feature of these five defective genes is their role in cellular processes the involve phosphorylation, particularly by members of the family of tyrosine kinases. For example insulin-like growth factor 1 receptor (**IGF1R**) belongs to the large family of tyrosine kinase receptors and is activated by a hormone called insulin-like growth factor 1 (IGF-1) and by a related hormone called IGF-2 [46]. SOM_DTP_ nodes in meta-clades 10 through 15 that are associated with defective **IGF1R**, occur where chemosensitivity mainly for leukemia cells. **IGF1R** is often overexpressed by tumors and mediates proliferation and apoptosis protection [47] [48]. As noted earlier [21], evaluation of drug sensitivity for compounds targeting leukemia cells has prompted the emergence of **IGF1R** as a potential therapeutic target for the treatment of leukemia. Weisberg, et al. [49] report that **IGF1R** protein expression/activity was substantially increased in mutant RAS-expressing cells, and suppression of RAS led to decreases in **IGF1R**. Synergy between MEK and **IGF1R** inhibitors correlated with induction of apoptosis, inhibition of cell cycle progression, and decreased phospho-S6 and phospho-4E-BP1. They suggested that given the complexity of RAS signaling, it is likely that combinations of targeted agents will be more effective than single agents, inclusive of **IGF1R** inhibitors.

**Table IV.** Group **C** (meta-clades 10 through 15). Table lists the defective genes that satisfy Fisher’s exact significance (p<0.05) for each meta-clade in Group **C**. Row entries list the counts for each FDA and MOA entry for significant defective genes.

| IGF1R | ATM | NOTCH1 | CDC25A | BRAF | FDA | MOA |
| --- | --- | --- | --- | --- | --- | --- |
| 3 | 0 | 2 | 1 | 0 | Arsenic_trioxide | Apo |
| 0 | 0 | 1 | 4 | 0 | (+)-JQ1 | BRD |
| 3 | 0 | 3 | 0 | 0 | Belinostat | HDAC |
| 0 | 0 | 0 | 3 | 1 | Entinostat | HDAC |
| 5 | 0 | 1 | 1 | 0 | Vorinostat | HDAC |
| 0 | 0 | 0 | 2 | 1 | Ixazomib_citrate | PSM |
| 3 | 0 | 6 | 1 | 0 | Dromostanolone_Propionate | Ho |
| 2 | 0 | 1 | 0 | 0 | 3-Bromopyruvate_acid | Mito\|HK2 |
| 5 | 0 | 2 | 0 | 0 | Dimethylaminoparthenolide | NFkB |
| 3 | 0 | 2 | 0 | 0 | Imexon | Apo |
| 3 | 0 | 3 | 0 | 0 | Dacarbazine | A7\|Ho\|AlkAg\|anti-Ho |
| 1 | 0 | 1 | 0 | 0 | 8-Chloro-adenosine | AM |
| 4 | 1 | 0 | 0 | 0 | Buthionine_sulphoximine | AM |
| 14 | 2 | 0 | 2 | 0 | Fenretinide | Ds |
| 9 | 4 | 1 | 0 | 0 | Fludarabine | Ds\|AM\|Dr |
| 5 | 1 | 2 | 1 | 0 | Fostamatinib | Ds\|Rs\|PK:YK,SYK |
| 0 | 0 | 0 | 3 | 1 | Pazopanib | PK:YK,PDGFR,FGFR,KIT,VEGFR |
| 1 | 0 | 1 | 4 | 0 | Axitinib | PK:YK,PDGFR,VEGFR |
| 2 | 0 | 4 | 0 | 0 | Imatinib | BCR-ABL\|PK:YK |

The evolutionarily conserved **NOTCH** family of receptors regulates a myriad of fundamental cellular processes including development, tissue patterning, cell-fate determination, proliferation, differentiation and, cell death[50]. The crosstalk among **Notch** and other prominent molecules/signaling pathways includes DNA damage repair[51]. DDR is a complex protein kinase based-signaling pathway which is conducted by the members of the phosphoinositide 3-kinase-like kinase (PIKK) family, such as ataxia telangiectasia mutated (**ATM**). **NOTCH1** is a major oncogenic driver in T cell acute lymphoblastic leukemia[52]. **NOTCH1** siRNA can effectively inhibit the expression of **NOTCH1** gene, inhibit the proliferation of lung cancer A549 cells and increase the sensitivity to chemotherapeutic drugs[53]. Of specific interest is the intersection of defective **NOTCH1** and the projection for Imatinib. Aljedai et al. [54] explored the role of **NOTCH1** signaling in chronic myeloid leukemia cells to find cross-talk between **NOTCH1** and BCR-ABL. Their results revealed that Imatinib induced BCR-ABL inhibition results in upregulation of **NOTCH1** activity. In contrast inhibition of **NOTCH1** leads to hyperactivation of BCR-ABL. They proposed that the antagonistic relationship between **NOTCH1** and BCR-ABL in CML suggests a combined inhibition of **NOTCH1** and BCR-ABL may provide superior clinical response over tyrosine-kinase inhibitor monotherapy.

Activation of checkpoint arrest and homologous DNA repair are necessary for maintenance of genomic integrity during DNA replication[55]. Germ-line mutations of the ataxia telangiectasia mutated (**ATM**) gene result in the well-characterized ataxia telangiectasia syndrome, which manifests with an increased cancer predisposition. Somatic **ATM** mutations or deletions are commonly found in lymphoid malignancies. Such mutations may be exploited by existing or emerging targeted therapies that produce synthetic lethal states. Cancers with mutations in genes encoding proteins involved in DNA repair may be more sensitive to treatments that induce synthetic lethality by inducing DNA damage or inhibiting complementary DNA repair mechanisms.

**CDC25A** affects cell proliferation and its expression is thought to be controlled through the PI3K-Akt-mTOR signaling pathway[56]. Sadeghi et al. [57] suggest that **CDC25A** controls the cell proliferation and tumorigenesis by a change in expression of proteins involved in cyclin D1 regulation and G1/S transition. The finding that defective **CDC25A** is associated with MOA:PK is consistent with the appearance of Pazopanib and Axitinib in the FDA compounds listed in **Table IV**.

**BRAF** is the least frequent of this set of five defective genes. Its association with Imatinib (**Table IV**) represents an unusual occurrence. Defective **BRAF** will be discussed in greater detail in the <u>RESULTS</u> meta-clade group **E** (meta clades 19 through 20). A distinguishing feature of its appearance in meta-clade 10 through 15 involves its co-existence with defective **NOTCH1**. As noted above there appears to be significant cross talk between BCR-ABL and **NOTCH1**.

### <u>RESULTS</u> Group D (meta-clades 16 through 18)

There are thirteen MOA types and 18 defective genes associated with meta-clade 16 through 18 (cf. **Figure 12).** MOA:Tu dominates these results, while MOA:PK and MOA:HSP90 appear with the next highest frequencies. The most frequent defective and significant genes include **ALK, MYC, TNKS, RBPJL, FLT1, MMP9, MST1R and SPTBN1**. **Table V** lists the FDA compounds associated with these defective genes. Most of these defective genes are involved with the mitotic component of tumor cell proliferation. For example, **MYC** encodes a nuclear phosphoprotein that has been implicated in the regulation of cell proliferation and the development of human tumors [58] and is regarded as a major determinant of mitotic cell fate [59]. Inhibition of microtubule polymerization has been reported to block mitosis and induce cell death [60]. Conacci-Sorell et al. [61] report the expression of **MYC** results in the induction of the actin-bundling protein fascin, formation of filopodia, and plays a role in cell survival, autophagy, and motility. **MYC** also recruits acetyltransferases that modify cytoplasmic proteins, including α-tubulin. Marzo-Mas et al. [62] find the antiproliferative activity of colchicine to inhibit tubulin polymerization to be modulated by the downregulation of **c-MYC** expression. Alexandrova et al. [63] report that the N-terminal domain of **c-MYC** associates with alpha-tubulin and microtubules. Marzo-Mas et al. [62] also found that tubulin binding compounds were able to downregulate the expression of the VEGF, hTERT and **c-MYC** genes. Others [64] have proposed targeting oncogenic **MYC** as a strategy for cancer treatment, proposing the destruction of a microtubule-bound **MYC** reservoir during mitosis contributes to vincristine’s anti-cancer activity [65]. Collectively these results support a role of defective **MYC** in chemosensitivity to tubulin targeting agents.

**Table V.** Group **D** (meta-clades 16 through 18). Table lists the defective genes that satisfy Fisher’s exact significance (p<0.05) for each meta-clade in Group **D**. Row entries list the counts for each FDA and MOA entry for significant defective genes.

| MYC | TNKS | RBPJL | FLT1 | MMP9 | SPTBN1 | MST1R | ALK | FDA | MOA |
| --- | --- | --- | --- | --- | --- | --- | --- | --- | --- |
| 2 | 2 |  |  |  |  |  |  | Actinomycin_D | Db |
|  |  |  |  |  | 1 | 1 |  | Belinostat | HDAC |
|  | 1 |  |  |  | 1 | 2 | 1 | Abiraterone | Ho\|CYP17A1 |
| 2 | 2 | 2 |  |  |  |  |  | Alvespimycin | HSP90 |
|  | 1 |  | 1 | 2 | 1 | 1 |  | Tanespimycin | HSP90 |
| 1 | 1 | 1 | 1 |  |  |  |  | Geldanamycin_analog | HSP90 |
| 2 | 1 | 1 |  |  |  |  |  | Pipamperone | NonCan:Psy |
|  |  |  |  |  | 1 | 1 |  | LDK-378 | PK:ALK |
|  | 2 | 3 |  | 3 | 2 | 4 |  | Crizotinib | PK:YK,MET,ALK |
| 4 | 1 | 1 |  |  |  |  |  | Carfilzomib | PSM |
| 2 | 4 |  |  |  |  |  |  | Daunorubicin | T2 |
| 1 | 3 |  |  |  |  |  |  | Doxorubicin | T2 |
| 2 | 4 |  |  |  |  |  |  | Epirubicin | T2 |
|  | 1 | 1 | 2 | 4 | 4 | 4 | 1 | Benzimate | Tu\|Tu-frag-stab |
| 3 | 4 | 2 | 8 | 4 | 3 | 1 | 4 | Docetaxel | Tu\|Tu-frag-stab |
|  | 2 | 1 | 1 | 1 | 1 | 1 | 1 | Dolastatin_10 | Tu\|Tu-frag-stab |
| 3 | 3 | 2 | 1 |  |  |  |  | Eribulin_mesilate | Tu\|Tu-frag-stab |
| 7 | 5 | 4 | 3 | 2 | 2 |  |  | Paclitaxel | Tu\|Tu-frag-stab |
| 9 | 9 | 6 | 2 | 3 | 3 | 1 | 1 | Vinblastine | Tu\|Tu-frag-stab |
| 4 | 3 | 3 |  |  | 1 |  |  | Vincristine | Tu\|Tu-frag-stab |
| 8 | 2 | 2 | 3 | 1 | 1 |  |  | Vinorelbine | Tu\|Tu-frag-stab |

**Figure 12:**
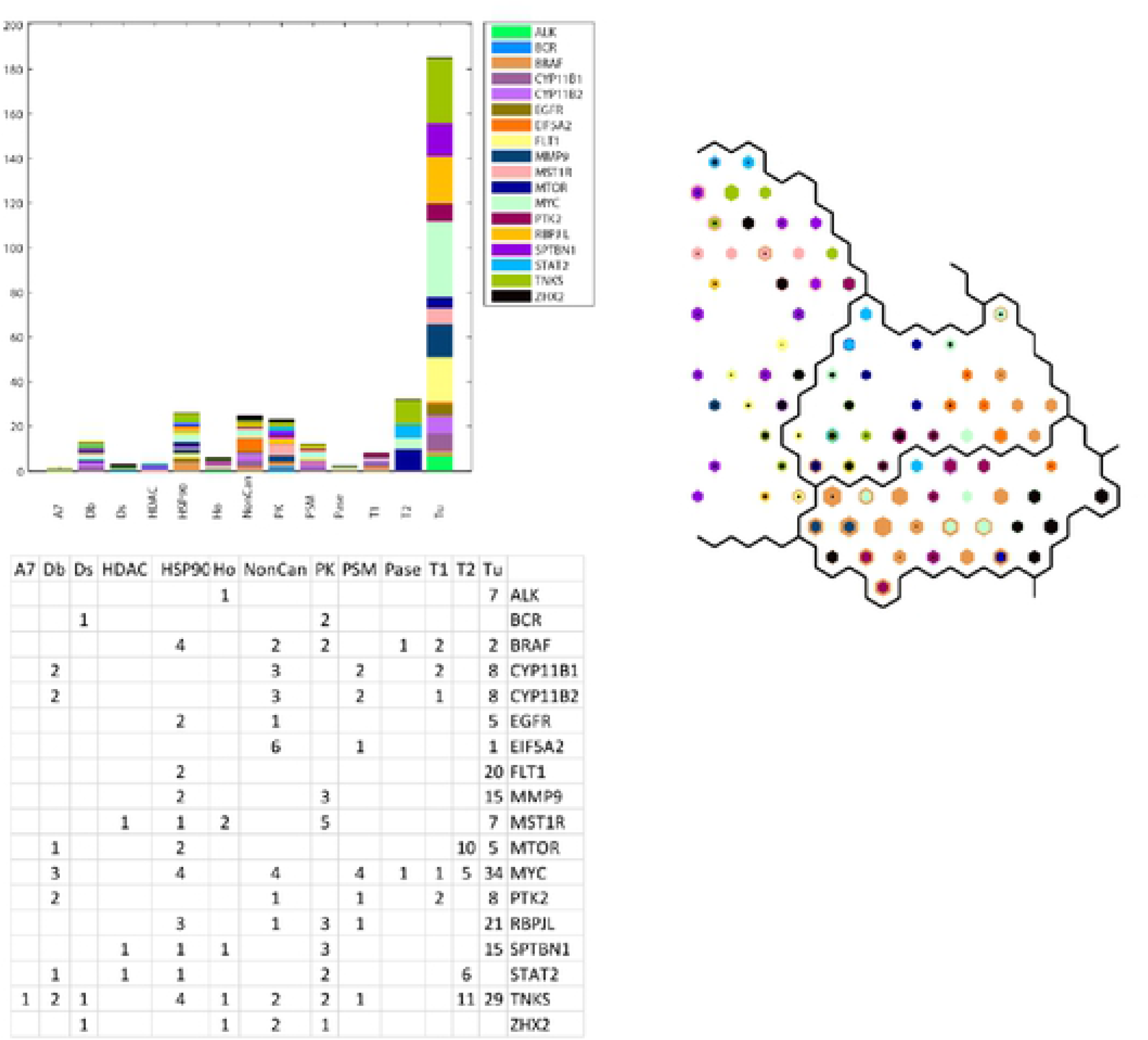
Upper left panel displays the histogram of counts for the co-occurrence of SOM_DTP_ nodes with defective genes and FDA compound projections. Lower left panel displays the counts for co-occurrence. Right panel displays the SOM_DTP_ for meta-clades 16 through 18 (Group **D**). See legend to **Figure 9** for additional details.

The second most frequent defective gene for MOA:Tu is **RBPJL. RBPJL** binds to DNA sequences almost identical to that bound by the Notch receptor signaling pathway transcription factor recombining binding protein J (**RBP-J**). A related family member RITA (RBPJ Interacting And Tubulin Associated 1) also acts as a negative regulator of the Notch signaling pathway that induces apoptosis and cell cycle arrest in human hepatocellular carcinoma [66]. Structural and biophysical studies demonstrate that RITA binds **RBP-J** and biochemical and cellular assays suggest that RITA interacts with additional regions on **RBP-J** [67]. Emerging evidence reveals Notch as a microtubule dynamics regulator and that activation of Notch signaling results in increased microtubule stability [68]. The **RBPJL**/RITA association raises the possibility that RITA-mediated regulation of Notch signaling may be influenced by **RBPJL** and potentially play a role in the chemosensitivity of Tu agents.

The third most frequent defective gene for MOA:Tu is **FLT1** (Fms-related tyrosine kinase (FLT) or VEGF receptor 1). The role of **FLT1** in the chemosensitivity of tubulin agents would appear to be unexpected. However, the blockade of VEGFR-1 and VEGFR-2 enhances paclitaxel sensitivity in gastric cancer cells [69]. Microtubule-targeted drugs inhibit VEGF Receptor-2 expression by both transcriptional and post-transcriptional mechanisms [70]. Novel anti-mitotics, which target the mitotic spindle through interactions with non-microtubule mitotic mediators like mitotic kinases and kinesins, have been identified and are now in clinical testing [71]. Included in clinical testing are compounds that have low nanomolar potency against ABL, **FLT1** and PDGFR [72]. Tumor endothelial cells demonstrate a strong activation of VEGF and Notch signaling [73]. VEGF-B is a growth factor that binds **FLT1** and is considered the odd member of the VEGF family, with mainly angiogenic and lymphangiogenic activities. VEGF-B has protective effects on neuropathy [74]. **FLT1** has been proposed as a prognostic indicator in endometrial carcinoma [75].

**MMP9** (matrix metalloproteinases 9) and its associated vascular endothelial growth factor (VEGF) are critical for tumor vascularization and invasion. A recent study of the expression of **MMP-9** and VEGF**(FLT1**) in breast cancer patients found their correlation significant enough to propose these genes as prognostic indicators [76]. Inspection of these SOM meta-clades finds MOA:PK agents to be located mainly in the upper portion of SOM meta-clade 16, where defective genes **MST1R**, **MMP9** and **RBPJL** also appear. Crizitonib is co-projected to these SOM nodes. Cizitonib is a small molecule TKI that inhibits the activity of the ALK fusion proteins, MET, ROS1, and **MST1R** (RON) [77, 78]. Defective MST1R may contribute to enhanced crizitonib chemosensitivity.

The defective **ALK** gene is the remaining gene listed for SOM meta-clades 16 through 18. Echinoderm microtubule-associated protein (EMAP)–like or EML proteins usually function in the cytoskeleton, contributing to the formation of the mitotic spindle and the interphase microtubule network. However, in around 5% of non–small cell lung cancers (NSCLC), chromosome rearrangements generate oncogenic fusion proteins that link parts of EML4 to the anaplastic lymphoma kinase (**ALK**). EML4–ALK proteins are highly transforming and pathogenic in NSCLC due to increased oligomerization and constitutive, kinase-activating autophosphorylation. However, at the same time, these fusion proteins create the therapeutic Achilles heel of an oncogenic addiction state in which the cancer cells depend on the tyrosine kinase activity of the fusion protein for survival—thus explaining the impressive clinical responses to crizotinib and other small-molecule drugs inhibiting the **ALK** tyrosine kinase [79].

MOA:HSP90 is the second most frequent MOA class in SOM meta-clades 16 through 18. Several studies have suggested a possible connection between HSP90 and the microtubule cytoskeleton. Weis et al. [80] find that HSP90 protects tubulin against thermal denaturation. Anti-tumor selectivity of a novel Tubulin and HSP90 dual-targeting inhibitor has been identified in non-small cell lung cancer model [81]. The presence of geldanamycin within the list of agents in this SOM region is consistent with this observation. Liu et al. ([82]) find evidence that mis-regulated HSP90 can affect drug sensitivity, an effect proposed to be due the altered regulation of HSP90 client proteins, inclusive of tubulin.

### <u>RESULTS</u> Group E (meta-clades 19 through 20)

Eleven MOA classes are found in SOM meta-clades 19 through 20 (cf. **Figure 13**). **BRAF** is the most frequently occurring defective gene in MOA:BCR-ABL and MOA:PK MOA, followed by **CDC25A, MET, BLM**, **RPTOR** and **PIK3CG** (cf. **Table VI**). This SOM_DTP_ region corresponds to the projection of known FDA approved **BRAF** targeting agents; dabrafenib, hypomethicin, selmutinub and vemurafenib. These results are consistent with the findings of Ikediobi et al. [4].

**Table VI.**
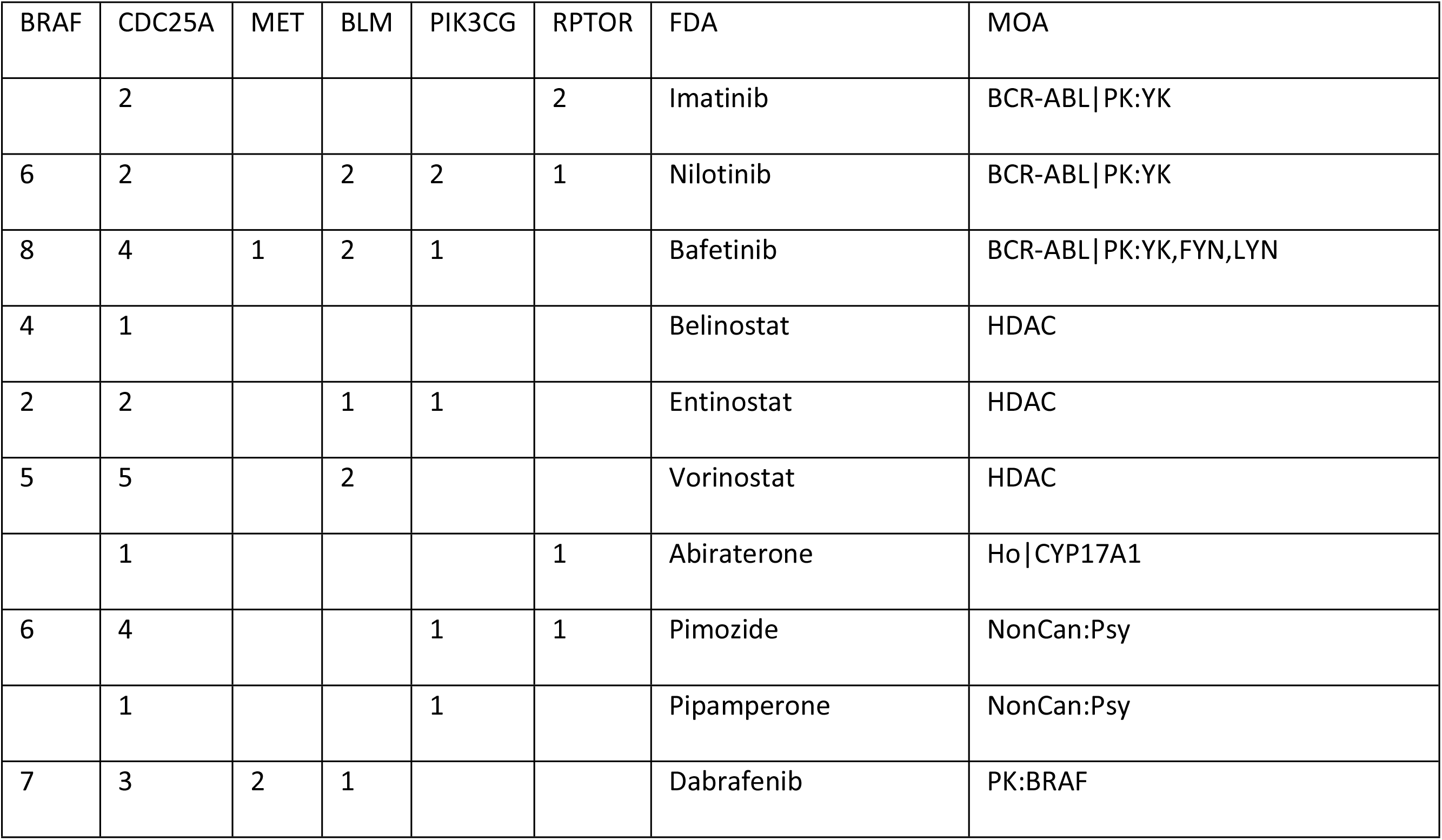
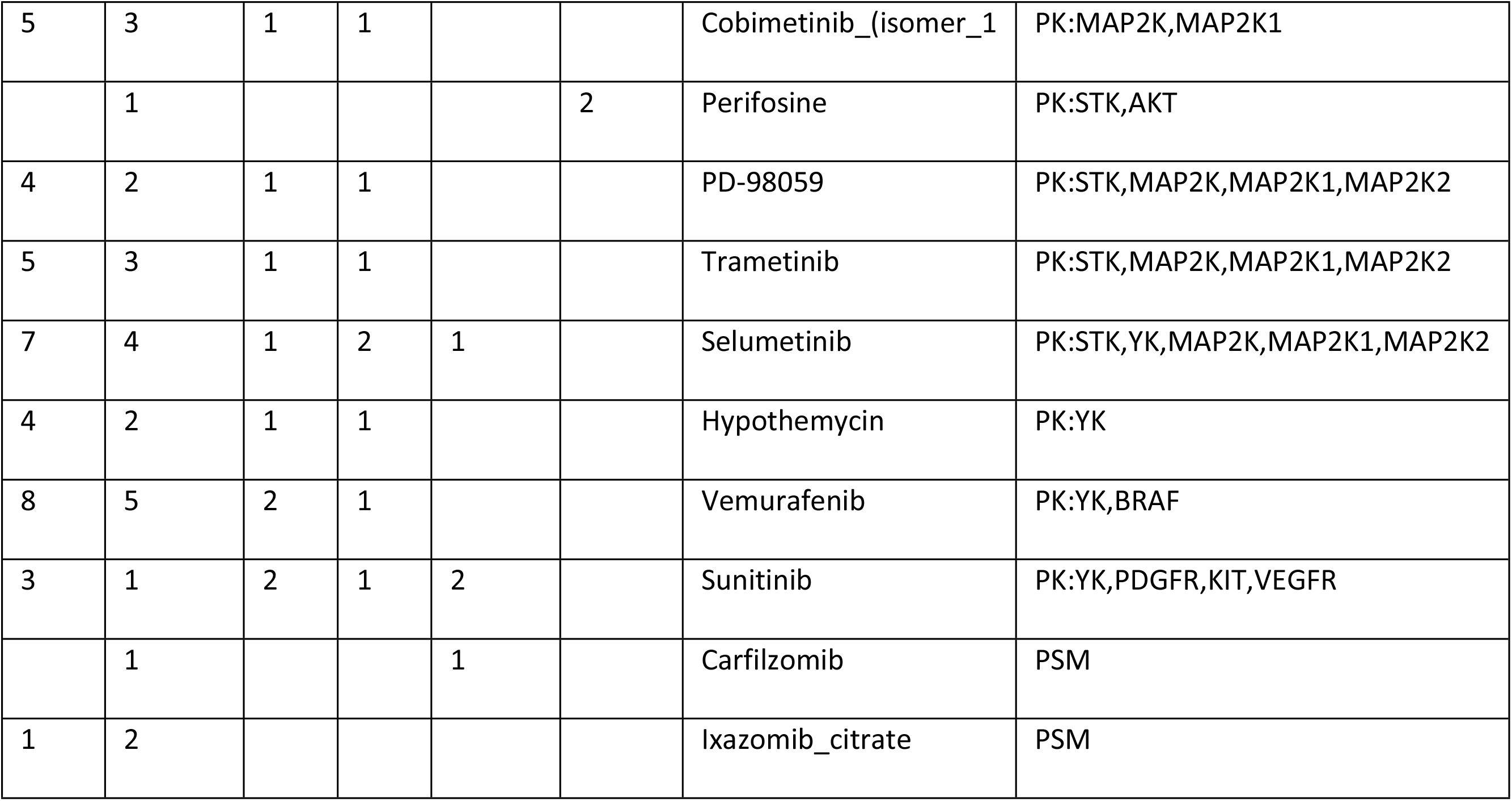
Group **E** (meta-clades 19 through 20). Table lists the defective genes that satisfy Fisher’s exact significance (p<0.05) for each meta-clade in Group **E**. Row entries list the counts for each FDA and MOA entry for significant defective genes.

| BRAF | CDC25A | MET | BLM | PIK3CG | RPTOR | FDA | MOA |
| --- | --- | --- | --- | --- | --- | --- | --- |
|  | 2 |  |  |  | 2 | Imatinib | BCR-ABL\|PK:YK |
| 6 | 2 |  | 2 | 2 | 1 | Nilotinib | BCR-ABL\|PK:YK |
| 8 | 4 | 1 | 2 | 1 |  | Bafetinib | BCR-ABL\|PK:YK,FYN,LYN |
| 4 | 1 |  |  |  |  | Belinostat | HDAC |
| 2 | 2 |  | 1 | 1 |  | Entinostat | HDAC |
| 5 | 5 |  | 2 |  |  | Vorinostat | HDAC |
|  | 1 |  |  |  | 1 | Abiraterone | Ho\|CYP17A1 |
| 6 | 4 |  |  | 1 | 1 | Pimozide | NonCan:Psy |
|  | 1 |  |  | 1 |  | Pipamperone | NonCan:Psy |
| 7 | 3 | 2 | 1 |  |  | Dabrafenib | PK:BRAF |
| 5 | 3 | 1 | 1 |  |  | Cobimetinib_(isomer_1 | PK:MAP2K,MAP2K1 |
|  | 1 |  |  |  | 2 | Perifosine | PK:STK,AKT |
| 4 | 2 | 1 | 1 |  |  | PD-98059 | PK:STK,MAP2K,MAP2K1,MAP2K2 |
| 5 | 3 | 1 | 1 |  |  | Trametinib | PK:STK,MAP2K,MAP2K1,MAP2K2 |
| 7 | 4 | 1 | 2 | 1 |  | Selumetinib | PK:STK,YK,MAP2K,MAP2K1,MAP2K2 |
| 4 | 2 | 1 | 1 |  |  | Hypothemycin | PK:YK |
| 8 | 5 | 2 | 1 |  |  | Vemurafenib | PK:YK,BRAF |
| 3 | 1 | 2 | 1 | 2 |  | Sunitinib | PK:YK,PDGFR,KIT,VEGFR |
|  | 1 |  |  | 1 |  | Carfilzomib | PSM |
| 1 | 2 |  |  |  |  | Ixazomib_citrate | PSM |

**Figure 13:**
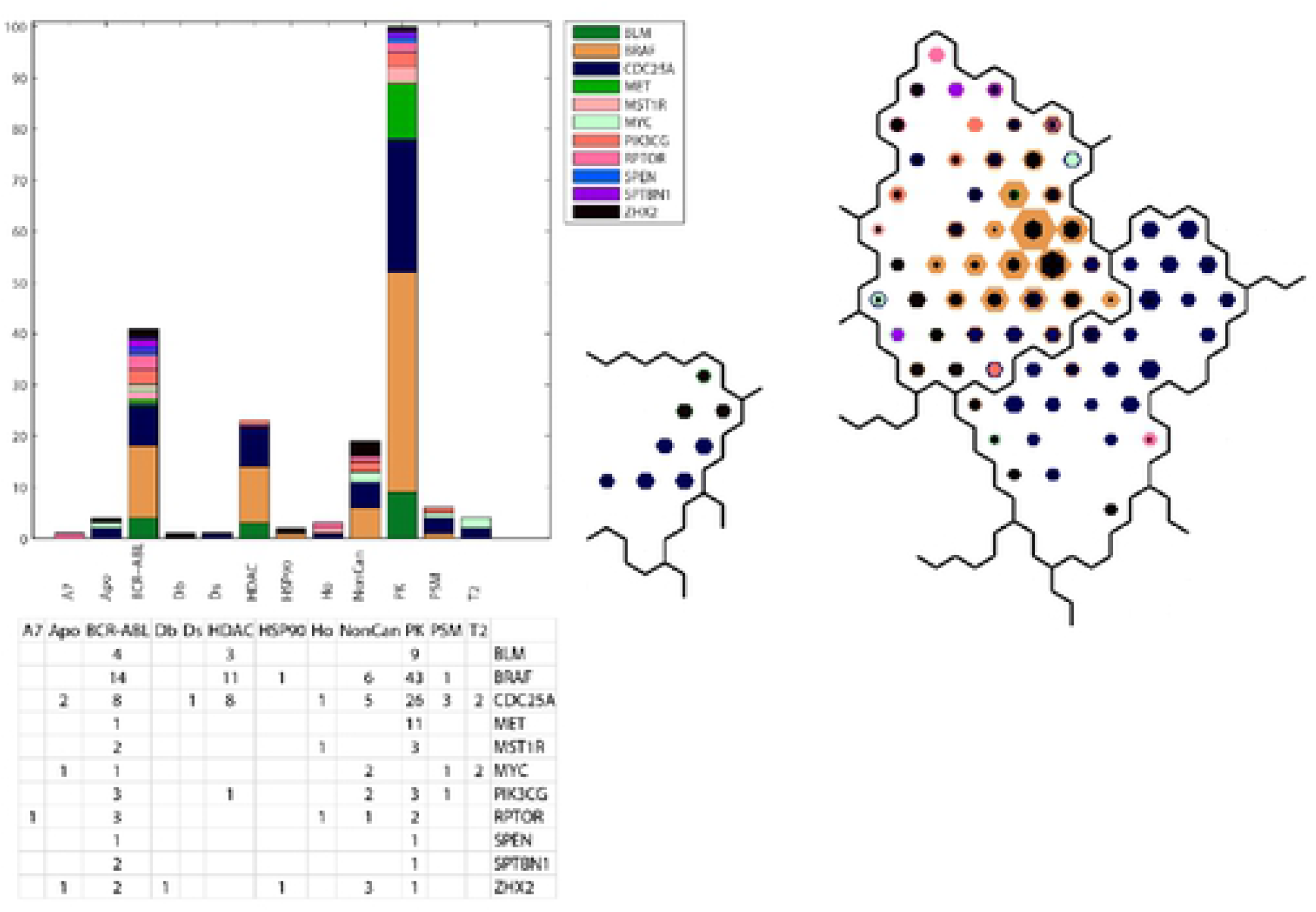
Upper left panel displays thehistogram of counts for the co occurrence of SOM_DTP_ nodes with defective genes and FDA compound projections. Lower left panel displays the counts for co-occurrence. Right panel displays the SOM_DTP_ for metacl ades 19 through 20 (Group E). See legend to **Figure 9** for additional details.

Defective **BRAF** is the most frequent defective gene in MOA:PK, BCR-ABL and HDAC. The association of defective **BRAF** with compounds that target this condition (**Table VI.**) are well documented[4, 83]. The association of defective BRAF to MOA:HDAC is consistent with literature reports. Mutant BRAF (v-Raf murine sarcoma viral oncogene homolog B1) inhibitors such as vemurafenib and dabrafenib have achieved unprecedented clinical responses in the treatment of melanomas[84, 85]. Recent studies have shown that histone deacetylase (HDAC) and mutant BRAF (v-Raf murine sarcoma viral oncogene homolog B1) inhibitors synergistically kill melanoma cells with activating mutations in BRAF by induction of necrosis[86].

Defective **CDC25A**, the second most frequent gene in MOA:PK, has been discussed in <u>RESULTS</u> Group **C** (meta-clades 10 through 15). The appearance of **CDC25A** in neighboring meta-clades 11 and 20 represents a case where meta-clade construction, based on hierarchical clustering, may have improperly grouped the 28 SOM_DTP_ meta-clades. Observation of **Figure 13** indicates that the projections of **BRAF** and **CDC25A** are into neighboring meta-clades (19 and 20). Thus, no association between **BRAF** and **CDC25A** is indicated.

Defective **MET** is the third most frequently occurring gene in the MOA:PK class. Virzi et al. [87] report the cancer carrying alterations of two oncogenes residing on the same pathway; namely, **MET** amplification and **BRAF** mutation. Surprisingly, the pharmacological blockade of **BRAF** had no effect, as it was followed by **MET** reactivation: Mechanistic studies unraveled the existence of a previously unknown negative feedback inhibition of **MET** by **BRAF**. **MET** inactivation in the context of the **BRAF**-activating mutation is driven through a negative feedback loop involving PP2A phosphatase, which in turn leads to phosphorylation of **MET** (Ser985). Disruption of this feedback loop allows PP2A reactivation, removing the inhibitory **MET** phosphorylation thereby unleashing **MET** kinase activity. These results provide an indication that **MET** targeting kinase agents may provide a dual therapeutic strategy for treating **BRAF** mutant tumors.

A role for defective **PIK3CG** is indicated in SOM meta clades 19 through 20 for MOA:PK. The publications from Shi et al. [88], Van Allen et al. [89] and Rizos et al. [90] addressed the roles of **PI3K** pathway gene’s mutations. Resistance to **BRAF** inhibitors can be associated with upregulation of the PI3K/AKT pathway, resulting from AKT1/3 mutations and mutations in positive (PIK3CA, **PIK3CG**) and negative (PIK3R2, PTEN and PHLPP1) regulatory genes.[91]

A role for defective **RPTOR** appears to be involved in MOA:BCR-ABL. Drugs simultaneously targeting two or more pathways essential for cancer growth could slow or prevent the development of resistant clones. Puausova et al. [92] identify dual inhibitors of proliferative pathways in human melanoma cells bearing the V600E activating mutation of **BRAF** kinase. They found these inhibitors to simultaneously disrupt the **BRAF** V600E-driven extracellular signal-regulated kinase (ERK) mitogen-activated protein kinase (MAPK) activity and the mechanistic target of rapamycin complex 1 (mTORC1) signaling in melanoma cells, yielding dynamic changes in mTOR(**RPTOR**) signaling.

The appearance of defective **BLM** (Bloom Syndrome RecQ Like Helicase), as potentially impacting MOAs BCR-ABL, PK and HDAC finds literature support. The Drosophila melanogaster ortholog of **BLM**, DmBlm, is encoded by mus309. Mutations in mus309 cause hypersensitivity to DNA-damaging agents and defects in repairing double-strand breaks [93].

The results in **Table VI** indicate possible role for HDAC in **BRAF** chemosensitivity. Gallagher et al. [94] find that HDAC inhibitors affect BRAF-inhibitor sensitivity by altering PI3K activity.

### <u>RESULTS</u> Group F (meta-clades 21 through 24)

The results for SOM meta-clades 21 through 24 appear in **Figure 14.** Eighteen MOA classes are found, with A7, Db, Df, Ds, T1 and T2 dominating this group. Eighteen defective genes exist, with **CDKN2A** as the most frequent defective gene, followed by **CDKN2B, AXL, IGF1R, MYC, STAT2, NRAS, KRAS, RPTOR and RB1**. Meta-clades 21 through 24 represent, by far, the largest number of FDA approved agents. Most of the most frequently occurring defective genes participate in DNA damage repair. However, none of these genes overlap with a prior analysis of DNA repair genes in the NCI60 and their predictive value for anticancer drug activity [95]. Su et al. [96] report that **CDKN2A** or **CDKN2B** loss is significantly associated with the sensitivity of CDK4/6 inhibitors (also projected to SOM meta-clade 14). Recent evidence indicates that **CDKN2A** is also involved in cell cycle independent functions such as DNA damage repair [97]. **CDKN2A** also provides instructions for making several proteins, including p16(INK4A) and p14(ARF), which function as tumor suppressors that keep cells from growing and dividing too rapidly or in an uncontrolled way. Overexpression of **CDKN2A** inhibits cell proliferation and invasion, and to cause cell cycle arrest in the G1 phase. **CDKN2A** mediates the AKT–mTOR (**RPTOR**) signaling pathway by suppressing lactate dehydrogenase (LDHA). [98]. Taken together, these results suggest therapeutic agents that target **CDKN2A** and **RPTOR i** n cancers that share these defective genes. Consistent with chemosensitivity for FDA compounds in these meta-clades, recent observations report that long term survivorship after high dose DNA damaging chemotherapy with melphalan is compatible with an increased chemosensitivity due to impairment of the DNA repair pathway [99]. Loss of CDK2 presents a different challenge to cells, aside from the more conventional role to regulate cyclins, which in turn might lead to altered DNA damage response and checkpoint activation, mutations in DNA repair genes drive cancer development [100, 101]. Noteworthy is the appearance of defective **RB1** (RB Transcriptional Corepressor 1) associated mainly with FDA compounds in MOA:A7. Although widely accepted as a key regulator of cell cycle progression by controlling the G1/S phase transition [1], recent evidence finds **RB1** to play a central role in DNA double strand break (DSB) repair[102]. **RB1** binds to tumor protein p53 binding protein 1 (53BP1) that directly links **RB1** function to the DNA damage response. Mutations of p53 and **RB1** are common in cancer cells [103]. Inactivation of **CDKN2A** can lead to deregulation of pathways containing these two defective genes.

**Figure 14.**
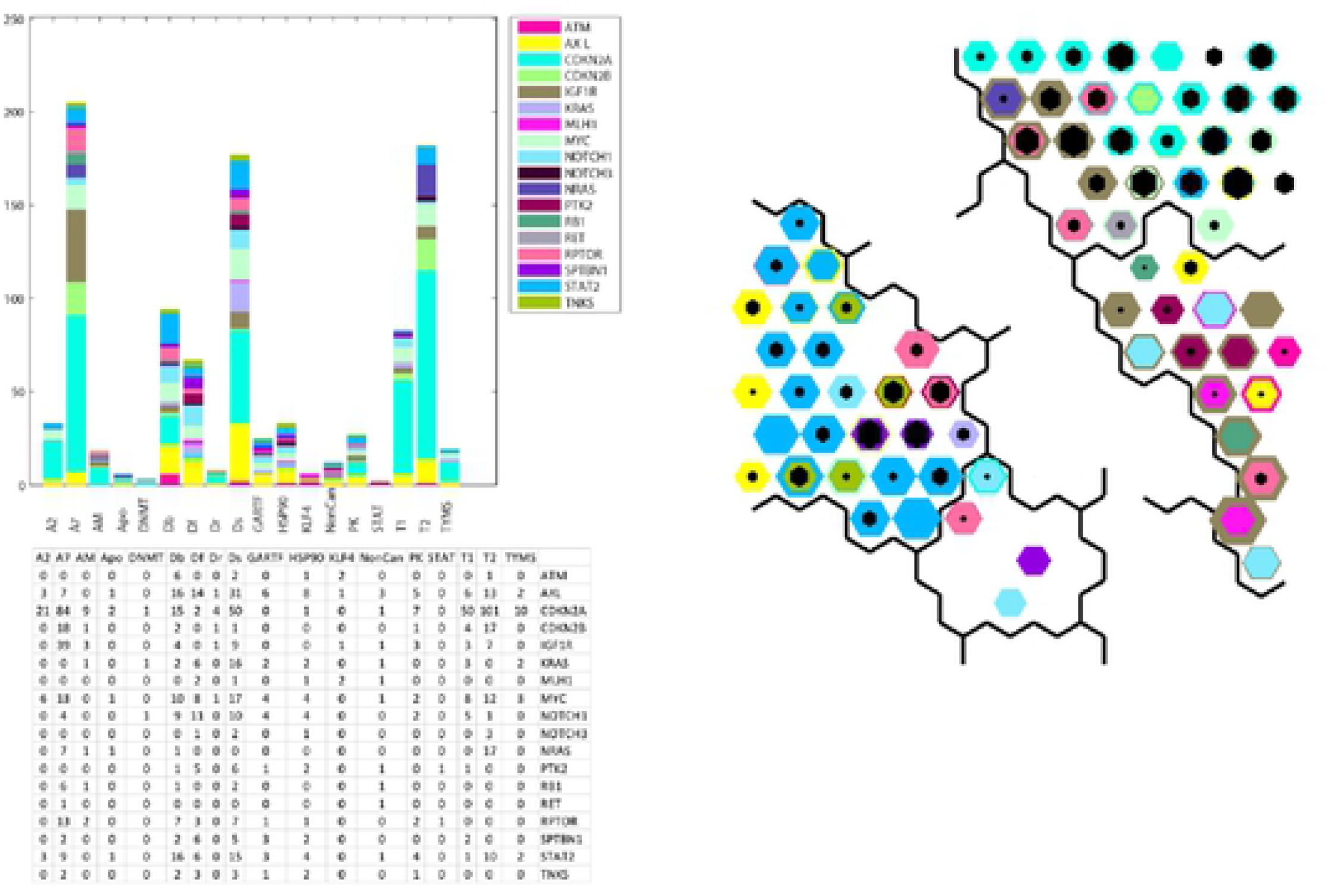
Upper left panel displays the histogram of counts for the co-occurrence of SOM_DTP_ nodes with defective genes and FDA compound projections. Lowerleft panel displays the counts for co occurrence. Right panel displays the SOM_DTP_ for meta clades 21 through 24 (Group F). See legend to **Figure 9** for additional details.

Ras proteins play a crucial role as a central component of the cellular networks controlling a variety of signaling pathways that regulate growth, proliferation, survival, differentiation, adhesion, cytoskeletal rearrangements and motility of a cell [104]. HRAS and **NRAS** also regulate the DNA damage response [105]. Mutant RAS-driven tumorigenesis arises independently of wild-type RAS isoforms, but recent evidence indicates wild-type isoforms are involved. Grabocka and colleagues [106] report how the loss of wild-type RAS alters oncogenic signaling and dampens the DNA-damage response, thereby affecting tumor progression and chemosensitivity. Since the MOA agents listed for SOM meta-clades 21 through 24 have roles in DNA damage, defective **CDKN2A** and **NRAS** may contribute to chemosensitivity of tumor cells to these agents. While targeting defective **KRAS** remains elusive [107], small molecule inhibitors are in the pipeline [108]. Exploration of NCI60 screened compounds that project to meta-clades 21 through 24 may provide a starting point for lead discovery.

Pyrazoloacridine, Palbociclib, Methotrexate, Fluorouracil, 8-Chloro-adenosine, Pralatrexate, Pemetrexed, Pelitrexol, By-Product_of_CUDC-305, 6-Mercaptopurine, Oxaliplatin appear most frequently in this SOM region. A study of gastric cancer patients detected a high frequency of mutations in MLL4, ERBB3, FBXW7, MLL3, mtor**(RPTOR)**, NOTCH1, PIK3CA, **KRAS**, ERBB4 and EGFR [109]. **KRAS** mutations have been reported as predictors of the response of lung adenocarcinoma patients receiving platinum-based chemotherapy [110] [111]. **NOTCH1** mutations target **KRAS** mutant CD8+ cells to contribute to their leukemogenic transformation [112] [113]. Notable in the list of FDA approved agents associated with SOM meta-clades 21 through 24 is oxaliplatin. Oxaliplatin-based chemotherapy is more beneficial in **KRAS** mutant than in **KRAS** wild-type metastatic colorectal cancer [114]. SOM meta-clade 21 is the location of cytarabine (ara-C) and is consistent with the conclusion of Ahmad et al [18] that adult AML patients carrying defective **KRAS** benefit from higher ara-C doses more than wt **KRAS** patients. Enhanced chemosensitivity of tumor cells with defective **KRAS** may represent a link to these observations.

**Table VII.**
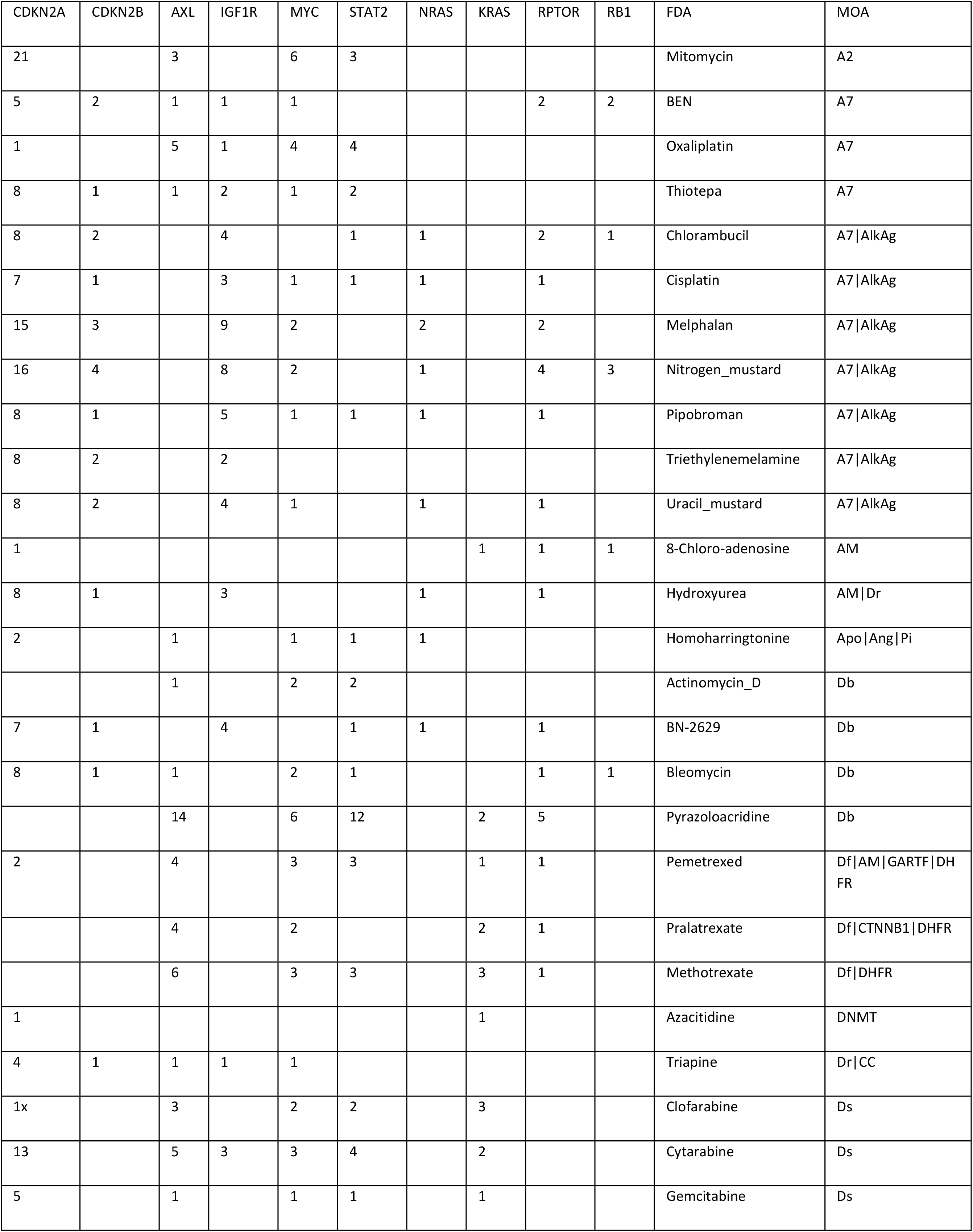
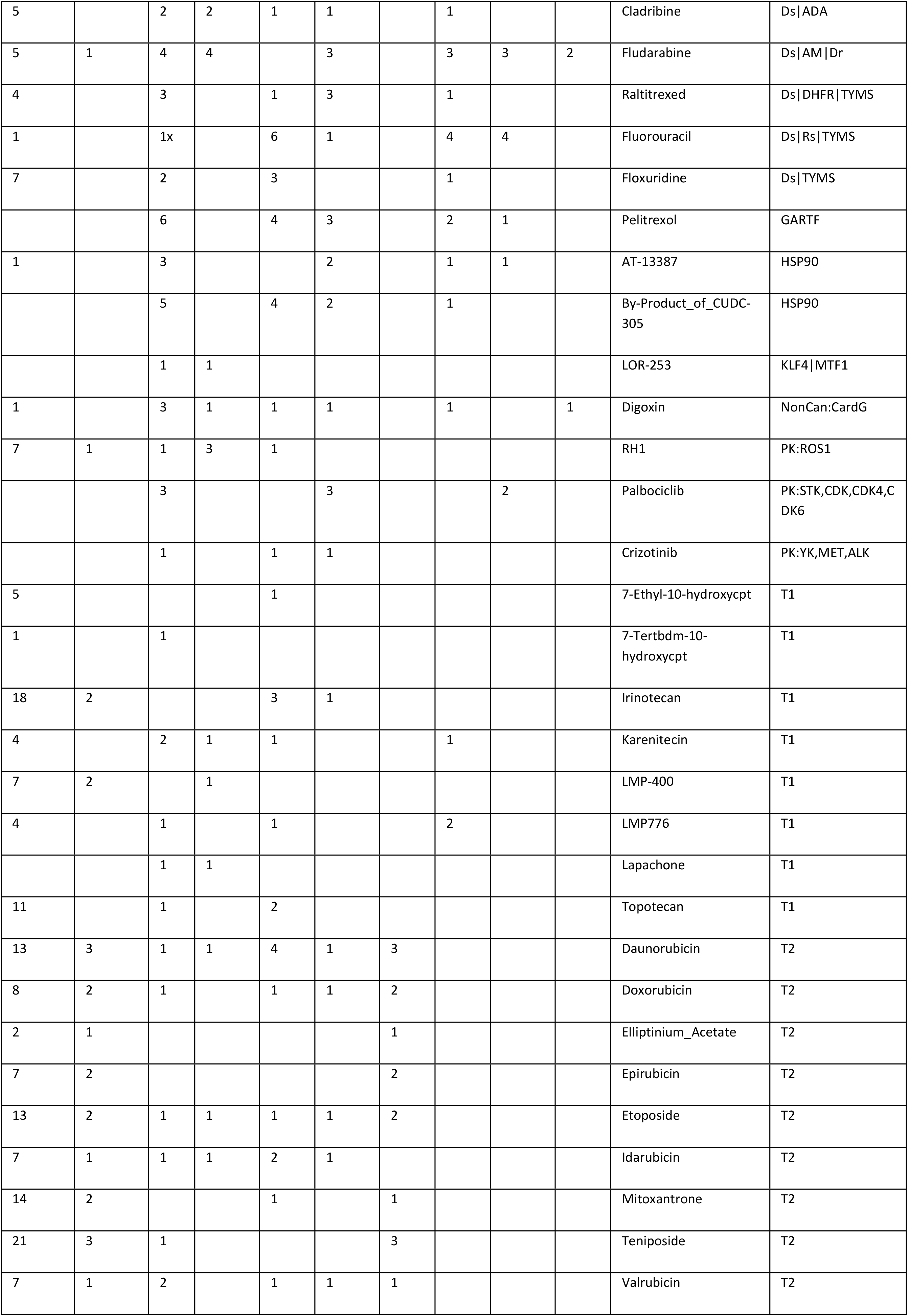
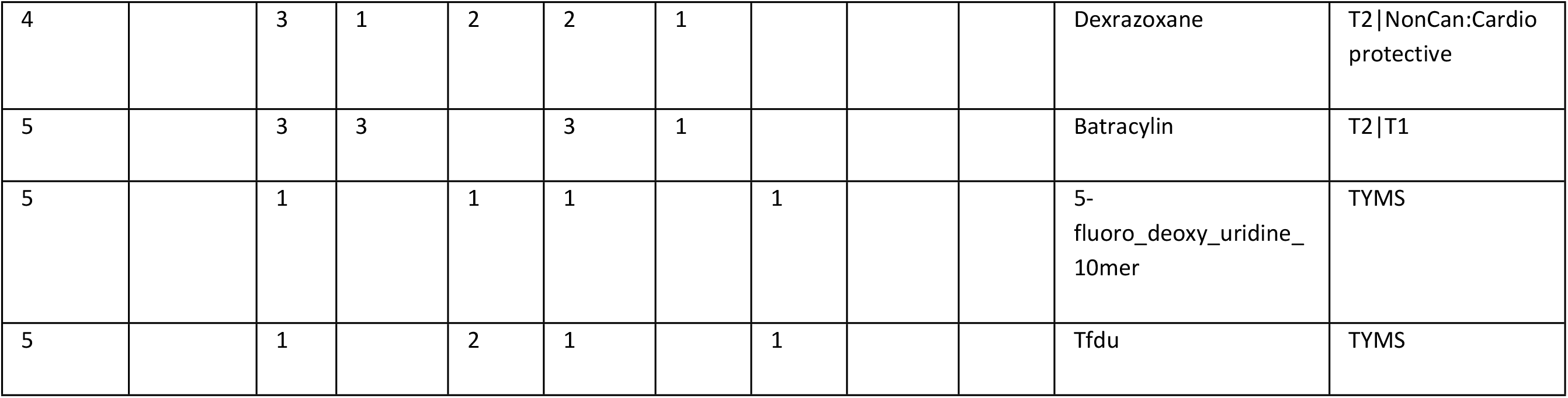
Group **F** (meta-clades 21 through 24). Table lists the defective genes that satisfy Fisher’s exact significance (p<0.05) for each meta-clade in Group **F**. Row entries list the counts for each FDA and MOA entry for significant defective genes.

### <u>RESULTS</u> Group G (meta-clades 24 through 28)

Eleven defective genes and fifteen MOA classes appear in meta-clades 24 through 28 (cf. **Figure 15**). The top-most frequently occurring defective genes are **RPTOR, NRAS, AXL, NOTCH1, MECOM**, **EGFR and EP300 (Table Next_8).** The defective genes in meta-clade 24 through 28 represent an amalgamation of the previous meta-clades, where sets of defective genes were involved in cellular processes of phosphorylation and progression through the cell cycle for proliferation. Consequently, all of the most frequent defective genes have been previously discussed, with the exceptions of **EGFR** and **EP300**.

**Figure 15.**
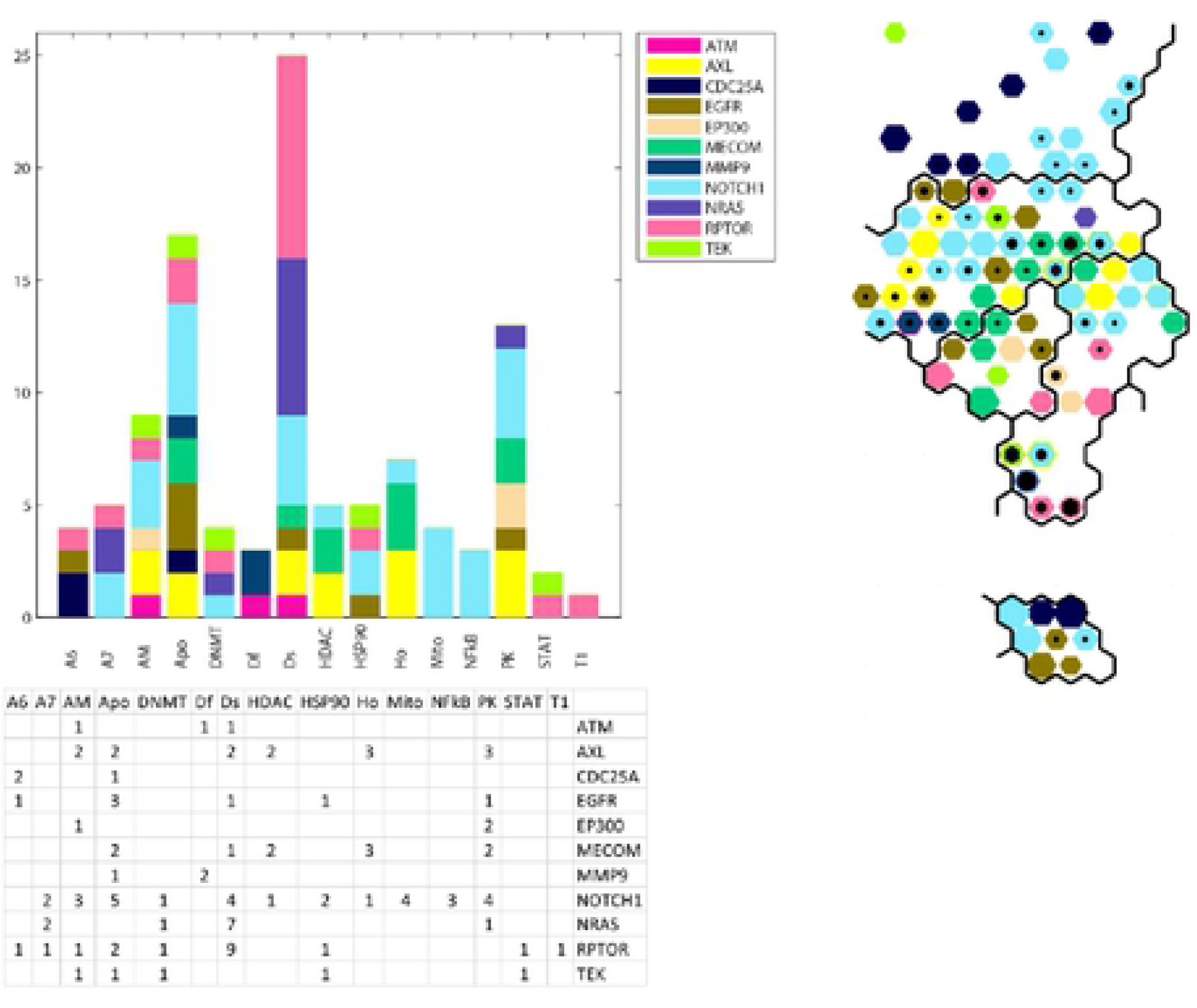
Upper left panel displays the histogram of counts for the co-occurrence of SOM_DTP_ nodes with defective genes and FDA compound projections. Lower left panel displays the counts for co occurrence. Right panel displays the SOM_DTP_ for meta clades 24 through 28 (Group G). See legend to **Figure 9** for additional details.

Epidermal growth factor receptor (**EGFR**) and its mutations contribute to tumorigenesis and biology of human cancers by affecting proliferation, progression and the process of apoptosis[115], with specific emphasis on defective **EGFR** and cell apoptosis[116]. Studies of mutant **EGFR** revealed that apoptosis was not prevented by inhibiting **EGFR** downstream proteins (PI3K, AKT and mTOR). These results identified another **EGFR** function, independent of PI3K/AKT/mTOR, which could be employed in therapy and drug design[115].

**EGFR** and **EP300**, as well as **NOTCH1**, are members of the ‘GO NEGATIVE REGULATION OF CELL CYCLE’ pathway, that stops, prevents or reduces the rate or extent of progression through the cell cycle. Defects in these genes could represent a cellular state whereby their reduced progression through the cell cycle could be perceived as drug-related growth inhibition.

Many tumors contained mutations in genes that regulate the cell cycle (TP53, CCND1, CDKN2A, FBXW7); epigenetic processes (MLL2, EP300, CREBBP, TET2); and the NOTCH (NOTCH1, NOTCH3), WNT (FAT1, YAP1, AJUBA) and receptor-tyrosine kinase-phosphoinositide 3-kinase signaling pathways (PIK3CA, EGFR, ERBB2).

**Table VIII.** Group **G** (meta-clades 25 through 28). Table lists the defective genes that satisfy Fisher’s exact significance (p<0.05) for each meta-clade in Group **G**. Row entries list the counts for each FDA and MOA entry for significant defective genes.

| RPTOR | NRAS | AXL | NOTCH1 | MECOM | EGFR | EP300 | FDA | MOA |
| --- | --- | --- | --- | --- | --- | --- | --- | --- |
| 1 |  |  |  |  | 1 |  | Lomustine | A6\|AlkAg |
| 1 | 2 |  | 1 |  |  |  | Dacarbazine | A7\|Ho\|AlkAg\|anti-Ho |
| 1 |  | 1 | 1 |  |  | 1 | 8-Chloro-adenosine | AM |
|  |  | 1 | 2 |  |  |  | Decitabine | AM\|DNMT |
| 1 |  | 1 | 2 | 2 | 1 |  | Curcumin | Apo |
| 1 |  | 1 | 1 |  | 2 |  | Obatoclax | Apo\|BCL2 |
| 1 | 1 |  | 1 |  |  |  | Azacitidine | DNMT |
|  |  |  |  | 1 | 1 |  | Cordycepin | Ds |
| 1 | 1 |  |  |  |  |  | Raltitrexed | Ds\|DHFR\|TYMS |
| 7 | 6 | 1 | 4 |  |  |  | 6-Mercaptopurine | Ds\|IMPDH2\|PPAT |
|  |  | 2 | 1 | 2 |  |  | Belinostat | HDAC |
|  |  | 3 | 1 | 2 |  |  | Raloxifene | Ho\|SERM |
| 1 |  |  | 1 |  |  |  | AT-13387 | HSP90 |
|  |  | 1 | 1 |  |  |  | 7-Hydroxystaurosporine | PK:STK |
|  | 1 | 1 | 2 |  |  |  | Triciribine_phosphate | PK:STK |
|  |  | 1 | 1 | 2 | 1 | 1 | Palbociclib | PK:STK,CDK,CDK4,CDK6 |

## DISCUSSION

The development of rational strategies for targeted cancer therapy will require integrative analysis of data derived from diverse sources including, but not exclusive to, large-scale, publicly available pre-clinical and clinical small-molecule screening and genomic data. A widely accepted difficulty towards linking screening and genomic data is how to gain molecular insight into the MOA(s) of active compounds. Not unexpectedly, the range of potentially important links is enormous; yielding massive challenges to the development of statistical/computational tools that assist integrative analyses. Advances have been made by focusing studies on fewer compounds (24 compounds in the CCLE [117] or approved FDA compounds [118]) or by studying small numbers of driver or mutated genes [119].

Following this strategy, the results of the present study demonstrate the power of combining genomic data and small-molecule screens of FDA compounds in the NCI60 to provide mechanistic clues about compound activity. These results reveal coarse-grained associations between chemosensitivity of target-directed FDA agents towards tumor cells harboring specific genetic defects. SOM clustering finds seven regions of <u>GI50</u>_NCI60_ responses, broadly assigned to FDA MOA classes that target Tubulin, BRAF mutations, RAF/MEK/ERK/mTOR and the PI3K/AKT pathways, DNA or protein synthesis pathways and are associated with a relatively unique set of defective genes for each MOA class. Salient associations include the role of defective MYC for tubulin targeting agents, defective CDKN2A, NRAS and KRAS for DNA damaging agents and the role of defective NOTCH1 for mutant BRAF targeting agents. Remarkably, nearly half of the defective genes reported herein also appear in Ikediobi et al.[4], albeit using very different methods.

Important caveats underly the interpretation of these results. <u>First</u>, links of defective genes to chemosensitivity are not revealed in a clear-cut manner. Rarely is chemosensitivity associated only to tumor cells harboring defective genes. Chemosensitivity also exists within tumor cells lacking defective genes (cf. **Figure 5**). Consequently, while gene-drug associations may provide a genetic basis for drug selection [120], there is clear evidence herein that additional, not well understood, factors are in play. <u>Second</u>, combinations of defective genes appear to play a role in chemosensitivity. For example, thirty-eight defective genes are listed in the Tables provided in each of the <u>RESULTS</u> subsections. Twenty-five of these defective genes are listed once, while thirteen appear more than once. Consequently, identifying a single defective gene as responsible for chemosensitivity will be rare; combinations are more likely. <u>Third</u>, global analysis of modest to large scale genomic and screening data offers only one perspective. The genetic make-up of the NCI60 represents only a snapshot of data for a small number of tumor cells. The universal application of results derived from the NCI60 may be relevant only in the rare instance that another tumor cell matches the genetic makeup of any NCI60 cell. This does not, however, rule out analyses, parallel to that presented here, that jointly examine existing and new data. <u>Fourth</u>, the absence of defective TP53 in these results has not gone undetected. Most NCI60 tumor cells harbor defective TP53. As a result, establishing a statistically significant Student’s t-test fails mainly due to too few responses of tumor cells lacking defective TP53. Extending the data analysis to more tumor cells, lacking mutant TP53, may prove helpful. <u>Fifth</u>, the most intriguing result of this analysis is the apparent support for synthetically lethal defective gene pairs as contributing to chemosensitivity. Defective genes exist withing the NCI60 as doublets, triplets, quartets, etc., and a subset of these genes are associated with tumor cells that exhibit chemosensitivity. Chemosensitive SOM_DTP_ nodes associated with tumor cells having more than one defective gene are also associated with numerous screened compounds. Combinations of these compounds may offer opportunities for targeting more than one defective gene in parallel pathways.

In summary, the challenge of finding meaningful results within complex and noisy data has been proposed using contemporary data and state-ot-the-art statistical tools. This global analysis of multiple datasets, overlapping in their origins within the NCI60, has provided a unique perspective of defective gene to drug MOA associations.

## ACKNOWLEDGEMENT

I would like to thank Drs. Ruili Huang and John Beutler for their extremely helpful comments made in the preparation of this manuscript.

